# A novel protein complex that regulates active DNA demethylation in *Arabidopsis*

**DOI:** 10.1101/2020.02.21.958371

**Authors:** Pan Liu, Wen-Feng Nie, Xiansong Xiong, Yuhua Wang, Yuwei Jiang, Pei Huang, Xueqiang Lin, Guochen Qin, Huan Huang, Qingfeng Niu, Jiamu Du, Zhaobo Lang, Rosa Lozano-Duran, Jian-Kang Zhu

**Author notes:** These authors contributed equally (Pan Liu, Wen-Feng Nie).

## Abstract

Active DNA demethylation is critical for altering DNA methylation patterns and regulating gene expression. The 5-methylcytosine DNA glycosylase/lyase ROS1 initiates a base excision repair pathway for active DNA demethylation and is required for the prevention of DNA hypermethylation at thousands of genomic regions in *Arabidopsis*. How ROS1 is regulated and targeted to specific genomic regions is not well understood. Here, we report the discovery of an *Arabidopsis* protein complex that contains ROS1, regulates *ROS1* gene expression, and likely targets the ROS1 protein to specific genomic regions. ROS1 physically interacts with a WD40 domain protein (RWD40), which in turn interacts with a methyl-DNA binding protein (RMB1) as well as with a zinc finger and homeobox domain protein (RHD1). RMB1 binds to DNA that is methylated in any sequence context, and this binding is necessary for its function *in vivo*.Loss-of-function mutations in *RWD40, RMB1*, or *RHD1* cause DNA hypermethylation at several tested genomic regions independently of the known ROS1 regulator IDM1. Because the hypermethylated genomic regions include the DNA methylation monitoring sequence in the *ROS1* promoter, plants mutated in *RWD40, RMB1*, or *RHD1* show increased *ROS1* expression. Importantly, ROS1 binding to the *ROS1* promoter requires RWD40, RMB1, and RHD1, suggesting that this complex dictates ROS1 targeting to this locus. Our results demonstrate that ROS1 forms a protein complex with RWD40, RMB1, and RHD1, and that this novel complex regulates active DNA demethylation at several endogenous loci in *Arabidopsis*.

## Introduction

DNA methylation at the fifth position of the cytosine ring is important for gene regulation, transposon silencing, and imprinting in plants and many other eukaryotic organisms (Robertson 2005; Slotkin and Martienssen 2007; Law and Jacobsen 2010; He et al. 2011). DNA methylation occurs in different sequence contexts, i.e. CG, CHG, and CHH, where H represents A, C, or T. In plants, *de novo* DNA methylation is mediated by the RNA-directed DNA methylation (RdDM)pathway, which involves small interfering RNAs (siRNAs) and scaffold RNAs in addition to an array of proteins (Law and Jacobsen 2010; Zhang et al. 2018). In *Arabidopsis*, CG methylation is maintained by DNA METHYLTRANSFERASE 1 (MET1) (Finnegan and Dennis 1993), while CHG methylation is maintained by the plant-specific DNA methyltransferase CHROMOMETHYLASE 3 (CMT3) (Cao and Jacobsen 2002). Asymmetric CHH methylation is maintained by DOMAIN REARRANGED METHYLTRANSFERASE 2 (DRM2) through the RdDM pathway and by CHROMOMETHYLASE 2 (CMT2) (Haag and Pikaard 2011; Zemach et al.2013).

Specific DNA methylation patterns are tightly regulated by DNA methylation and demethylation pathways (Penterman et al. 2007; Hsieh et al. 2009; Zhu 2009; Law and Jacobsen 2010; Furner and Matzke 2011; Matzke and Mosher 2014). Passive DNA demethylation results from an absence of DNA methyltransferases or a reduction in their activity, or from a shortage of methyl donors following DNA replication (Zhang et al. 2018). In plants, active DNA demethylation is mediated by 5-methylcytosine DNA glycosylases through a DNA base-excision repair pathway (Choi et al. 2002; Gehring et al. 2006; Zhu et al. 2007; Zhu 2009). There are four 5-methylcytosine DNA glycosylases in *Arabidopsis*, namely REPRESSOR OF SILENCING 1 (ROS1), DEMETER (DME), DEMETER-LIKE 2 (DML2), and DEMETER-LIKE 3 (DML3) (Choi et al. 2002; Gong et al. 2002; Penterman et al. 2007; Hsieh et al. 2009). Because the 5-methylcytosine DNA glycosylases can recognize and directly remove the 5-mC base, they are critical enzymes in the active DNA demethylation pathway and are thus often referred to as DNA demethylases (Zhang et al. 2018). Research on how these DNA demethylases are regulated and recruited to their target loci is required to reveal how distinct genomic DNA methylation patterns are generated and modified during organismal development and in response to environmental change.

The histone acetyltransferase Increased DNA Methylation 1 (IDM1) is part of the IDM protein complex, which specifically recognizes certain methylated genomic regions through the Methyl-CpG-Binding domain (MBD) of its subunits Methyl-CpG-Binding protein MBD7 and IDM1, and through the DNA-binding domain of HARBINGER TRANSPOSON-DERIVED PROTEIN HDP2 (Qian et al. 2012; Qian et al. 2014; Lang et al. 2015; Duan et al. 2017). Recent research showed that ROS1 physically interacts with the histone variant H2A.Z and is recruited to some genomic regions via SWR1-mediated H2A.Z deposition, which involves specific histone acetylation marks created by the IDM complex (Nie et al. 2019). Because this mechanism only applies to a subset of target genomic regions of ROS1, there must be alternative mechanisms for the targeting of ROS1.

In this study, we found that ROS1 forms a protein complex with RWD40, RMB1, and RHD1, which are proteins that contain WD40, methyl-DNA binding, and homeodomain and zinc finger domains, respectively. We show that this novel protein complex, which we call the RWD40 complex, is required for the prevention of DNA hypermethylation at several genomic regions in *Arabidopsis*, and that this function is independent of the IDM-SWR1-H2A.Z pathway. One of these regions is the *ROS1* promoter, which contains a DNA methylation monitoring sequence (MEMS) important for DNA methylation homeostasis in *Arabidopsis* (Lei et al. 2015; Williams et al. 2015). Consistent with its role in controlling the DNA methylation of the *ROS1* promoter, we found that this protein complex negatively regulates the expression of *ROS1*; importantly, ROS1 binding to the MEMS requires RWD40, RMB1, and RHD1, suggesting that the RWD40 complex dictates ROS1 targeting to this locus. Moreover, our results indicate that this complex plays a role in anti-bacterial resistance, consistent with the known function of ROS1 in plant-bacteria interactions. Taken together, our results suggest that RWD40, RMB1, and RHD1 act in concert with ROS1 to control genomic DNA methylation by regulating *ROS1* gene expression and likely by dictating the targeting of the ROS1 protein to selected target regions.

## RESULTS

### A WD40 protein physically interacts with ROS1 and functions in DNA demethylation

To date, two proteins, H2A.Z and MET18 (homolog of yeast MET18, Methionine 18), have been reported to physically interact with ROS1 and to regulate active DNA demethylation in *Arabidopsis* plants (Duan et al. 2015; Nie et al. 2019). H2A.Z is required for ROS1 to target specific genomic regions defined by the IDM and SWR1 protein complexes (Qian et al. 2012; Lang et al. 2015; Nie et al. 2019). MET18 is a critical enzyme in the biosynthesis of the MoCo cofactor required for the methyl-DNA glycosylase/lyase activity of ROS1 (Duan et al. 2015). To identify additional ROS1-interacting proteins, we used yeast two-hybrid (Y2H) assays to determine whether ROS1 interacts with selected proteins found in the anti-FLAG pull-down products from *ROS1-3xFLAG-3xHA* (*ros1-13* mutant expressing *3xFLAG-3xHA*-tagged *ROS1* under its native promoter) *Arabidopsis* plants (Lei et al. 2015) in previous experiments. The results showed that a WD40 domain-containing protein (which we named RWD40 for <u>R</u>OS1-associated <u>WD40</u> domain-containing protein) interacts with ROS1 (Figure 1A and 1B). The interaction between ROS1 and RWD40 was confirmed by bimolecular fluorescence complementation (BiFC) (Figure 1C). Transient expression assays in *Nicotiana benthamiana* leaves showed that RWD40 is mainly localized in the nucleus (Figure S1A). In the anti-FLAG immunoprecipitate from *ROS1-3xFLAG-3xHA* plants (Lei et al. 2015), but not from Col-0 control plants, we identified multiple RWD40 peptides (Figure 1D). Furthermore, we also identified peptides corresponding to ROS1 in the anti-FLAG immunoprecipitate from transgenic plants expressing a *3xFLAG-RWD40* fusion driven by the *RWD40* native promoter in the *rwd40-1* mutant background (Figures1D and S1B-S1D). These results strongly suggest that ROS1 physically interacts with RWD40 in *Arabidopsis*.

**Figure 1.**
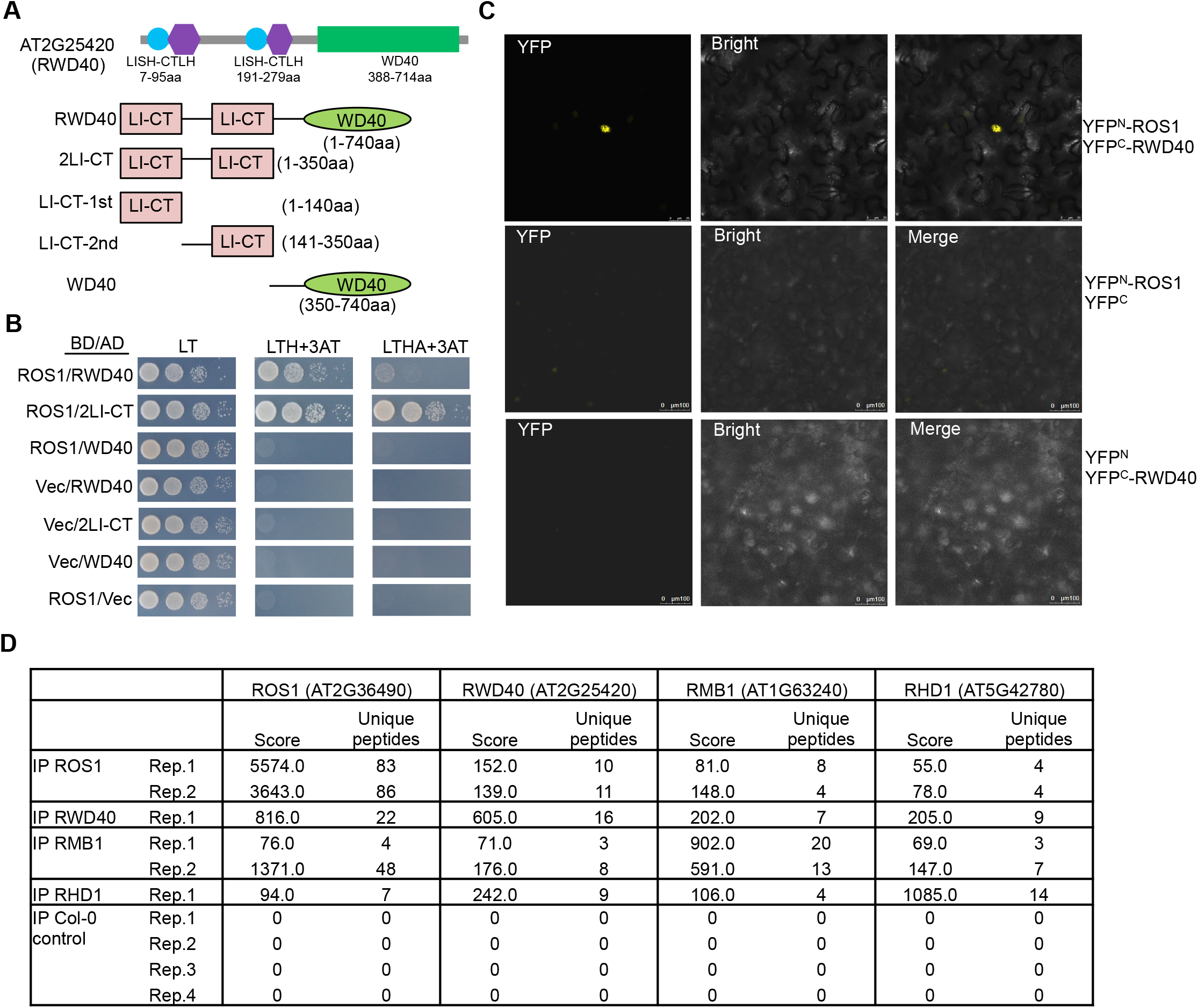
RWD40 interacts with ROS1. **(A)** Schematic representation of the RWD40 protein (top panel) and of the truncated forms of RWD40 used in Y2H assays (lower panels). The two LISH-CTLH-containing regions of RWD40 encompass amino acid residues 7 to 95 and 191 to 279, respectively. The WD40 domain encompasses amino acid residues 388 to 741. LI-CT indicates the LISH-CTLH domain. aa, amino acid residue. **(B)** RWD40 interacts with ROS1 in Y2H assays. The N-terminal region, including the two LISH-CTLH domains of RWD40, interacts with ROS1. **(C)** RWD40 interacts with ROS1 in BiFC assays. **(D)** Proteins detected by LC-MS/MS following immunoprecipitation (IP) using an anti-FLAG antibody, which immunopurifies ROS1, RWD40, RMB1, and RHD1 from transgenic Arabidopsis lines. Results obtained in one to four independent biological replicates are shown (Rep1–Rep4).

To investigate whether RWD40 regulates ROS1-dependent active DNA demethylation, we used Chop-PCR to determine the DNA methylation level at the AT5G39160 locus in *rwd40-1* and *rwd40-2* mutant seedlings. Similar to *ros1* mutant plants (Qian et al. 2012), *rwd40-1* and *rwd40-2* plants showed increased DNA methylation at the 5’ region of AT5G39160 (Figure S1E). Individual locus bisulfite sequencing confirmed that the DNA methylation levels at the AT5G39160 locus were higher in *rwd40-1, rwd40-2*, and *ros1-4* mutant plants than in the wild-type control (Figure S1F). Interestingly, the methylation level at AT5G39160 was not increased in the *idm1-2* mutant (Figure S1E and S1F). The expression of *RWD40-3xFLAG* fusion driven from its native promoter in the *rwd40-1* mutant reduced the methylation level at this locus (Figure S1E). These results show that RWD40 interacts with ROS1 and functions in regulating DNA demethylation.

### RMB1 and RHD1 interact with RWD40 and function in DNA demethylation

We noticed that the anti-FLAG immunoprecipitates from *3xFLAG-RWD40* transgenic plants contained peptides corresponding to AT1G63240 and AT5G42780, in addition to peptides from RWD40 and ROS1 (Figure 1D). AT1G63240 and AT5G42780 peptides were also detected in the anti-FLAG immunoprecipitates from *ROS1-3xFLAG-3xHA* transgenic plants (Figure 1D). The AT1G63240 protein is predicted to have a non-canonical methyl-DNA binding domain, a CBD domain, and a domain of unknown function, and is hereafter referred to as RMB1 (for <u>R</u>OS1-associated <u>m</u>ethyl-DNA <u>b</u>inding protein <u>1</u>) (Figure S2A). The AT5G42780 protein contains a zinc-finger domain and a homeodomain, and is hereafter referred to as RHD1 (for <u>R</u>OS1-associated <u>h</u>omeo<u>d</u>omain protein 1) (Figure S2B). In Y2H assays, RWD40 interacts with RMB1 and RHD1, and RMB1 interacts with RHD1 (Figure 2A). These interactions as well as the interaction between RWD40 and ROS1 were confirmed by split luciferase and BiFC assays in *N. benthamiana* leaves (Figures 2B and S2C). The assays also show that neither RMB1 nor RHD1 can directly interact with ROS1 (Figures 2A, 2B and S2C).

**Figure 2.**
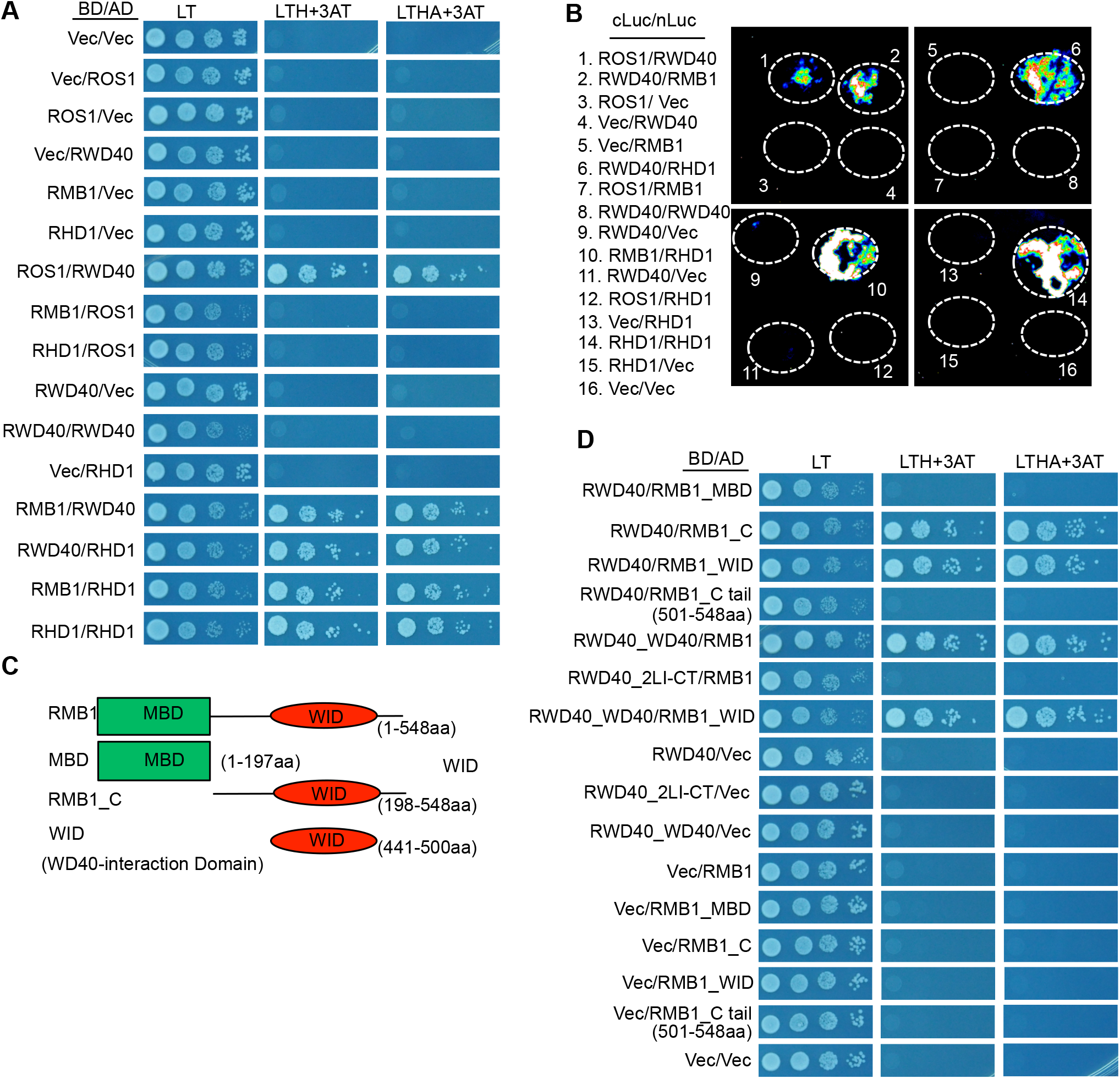
RMB1 and RHD1 interact with RWD40 but not with ROS1. **(A)** Interactions among RWD40, RMB1, RHD1, and ROS1 in Y2H assays. BD, GAL4-binding domain; AD, GAL4-activation domain. **(B)** Interactions among RWD40, RMB1, RHD1, and ROS1 in split luciferase complementation assays. The indicated proteins were fused to the N-terminal or the C-terminal part of the luciferase protein (nLuc or cLuc, respectively) and transiently expressed in *Nicotiana benthamiana* leaves. White circles indicate regions infiltrated with *Agrobacterium*. The luminescent signal indicates a protein–protein interaction. **(C)** Schematic representation of the truncated forms of RWD40 used in the Y2H assays. WID, WD40-interaction domain. aa, amino acid residue. **(D)** The WID domain of RMB1 interacts with the WD40 domain of RWD40 in Y2H assays.

RMB1 was immunoprecipitated from transgenic plants expressing a *3xFLAG-RMBD1* fusion driven by the RMB1 native promoter in the *rmb1-1* mutant background (Figure S2A). The immunoprecipitate contained not only RMB1 but also RWD40, RHD1, and ROS1 (Figure 1D). Similarly, the RHD1 immunoprecipitate from transgenic plants expressing a *3xFLAG-RHD1* fusion driven by the RHD1 native promoter in the *rhd1-1* background (Figure S2B) contained not only RHD1 but also RMB1, ROS1, and RWD40 (Figure 1D). These results show that RWD40, RMB1, and RHD1 proteins are associated with ROS1 *in vivo*.

Dysfunction of *RMB1* caused DNA hypermethylation at the AT5G39160 locus (Figures S2D and S2E). The DNA hypermethylation phenotype of the *rmb1-1* mutant was suppressed in transgenic plants expressing *3xFLAG-RMB1* (Figure S2D). The *rhd1-1* mutant, however, did not show a DNA hypermethylation phenotype at the AT5G39160 locus (Figure S2F). We assessed the methylation levels at another endogenous ROS1 target, AT4G18380, by Chop-PCR. Similar to *ros1-4* mutant plants, *rhd1-1* mutant plants had increased DNA methylation at AT4G18380 (Figure S2F). Expression of the *3xFLAG-RHD1* transgene suppressed this hypermethylation phenotype of the *rhd1-1* mutant (Figure S2F). These results suggest that RMB1 and RHD1 function in regulating locus-specific demethylation in *Arabidopsis*.

### RWD40, RMB1, RHD1, and ROS1 form a protein complex

The interactions of RWD40 with ROS1, RMB1, and RHD1, and between RMB1 and RHD1 in Y2H, BiFC, and split luciferase assays, together with their *in vivo* association as indicated by immunoprecipitation-mass spectrometry analyses, suggested that the four proteins may form a protein complex in plant cells. To assess this possibility, we carried out gel filtration assays with protein extracts from *3xFLAG-RMB1, 3xFLAG-RWD40, 3xFLAG-RHD1*, and *ROS1-3xFLAG-3xHA* transgenic plants. The results showed that RMB1, RWD40, RHD1, and ROS1 were eluted in the same fractions, and indicated that they exist in a complex with an estimated size of approximately 350 kDa (Figure 3A). This is close to the predicted total size of 328 kDa if the four proteins are in a 1:1:1:1 ratio. We co-expressed full-length RWD40, RMB1, and RHD1, and a truncated ROS1 protein (amino acid residues 1 to 100) in Sf9 insect cells using baculovirus vectors. After conducting histidine affinity purification and gel filtration, we found that the four proteins were eluted in a single peak fraction (Figure 3B). All the gel bands (Figure 3B) were extracted and their identities were confirmed by mass spectrometry (Table S1). Taken together, these results suggest that RWD40, RMB1, RHD1, and ROS1 form a protein complex *in vivo*, referred to as the RWD40 complex hereafter.

**Figure 3.**
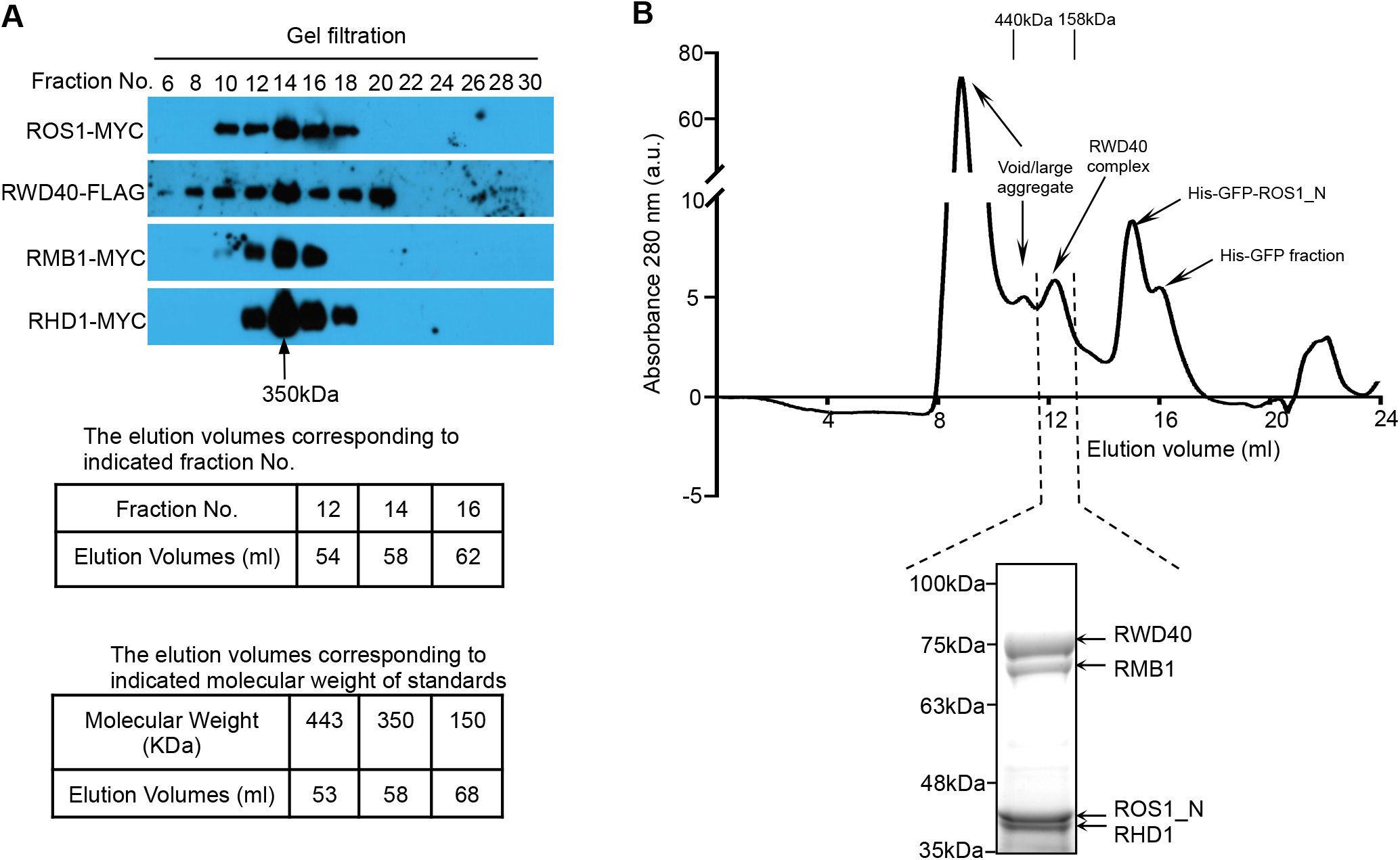
ROS1, RWD40, RMB1, and RHD1 form a protein complex. **(A)** Western blot analysis of gel filtration eluates. The indicated fractions eluted from the gel filtration column were probed with anti-MYC or anti-FLAG to detect the epitope-tagged RWD40, RMB1, RHD1, and ROS1. The arrow indicates the estimated molecular weight of the protein complex. A 120-ml column of HiPrep 16/60 Sephacryl S-300 HR was used. The elution volumes corresponding to the indicated fraction No. and indicated molecular weight of standards are shown. **(B)** Co-expression of full-length *RWD40, RMB1, RHD*, and a truncated *ROS1* protein (amino acid residues 1-100) in an insect cell expression system. RWD40, RMB1, RHD, and the truncated ROS1 were obtained in a single peak fraction. The positions of elution markers and their molecular masses (in kDa) are indicated. a.u., arbitrary units. A 24-ml column of Superdex200 10/300 GL was used for the Sf9 recombinant proteins.

### Identification of the protein domains that mediate the interactions within the protein complex

ROS1 contains an HhH-GPD domain, an End-III domain, a CXXC domain, an RRM_DME domain, and an N-terminal domain of unknown function (Figure S3A). The ROS1 N-terminal region (amino acid residues 41 to 100) has sequence similarity to the EAR (Ethylene response factor–associated <u>A</u>mphiphilic <u>R</u>epression) motif (Yang et al. 2018) (Figure S3A), and is thus referred to as the ELD (<u>E</u>AR-like <u>D</u>omain) domain. To determine which domain in ROS1 may mediate the interaction between ROS1 and RWD40, we performed Y2H assays with various deletion mutants of ROS1. We found that the N-terminal half (amino acid residues 1 to 696) of ROS1 interacts with RWD40, whereas the C-terminal half (amino acid residues 697 to 1393) does not (Figure S3B). Further truncation of the N-terminal half of ROS1 showed that the ELD domain (amino acid residues 41 to 100) is sufficient to mediate the interaction with RWD40 (Figure S3B).

We also performed Y2H assays with various deletion mutants of RWD40, and found that the N-terminal half of RWD40, including two LISH-CTLH domains but not the WD40 domain, is sufficient to enable the interaction with ROS1 (Figures 1B and S3B). On the other hand, the WD40 domain of RWD40 interacts with RMB1 in Y2H assays (Figure 2D). The C-terminal region of RMB1 interacts with the WD40 domain of RWD40 (Figures 2C and 2D), and is thus referred to as the WID (WD40 interaction domain) domain (Figure 2C). The WD40 domain of RWD40 is also capable of mediating the interaction between RWD40 and RHD1 (Figures S3C and S3D). The zinc-finger domain of RHD1 is responsible for the interaction with RWD40, while both the zinc finger domain and homeodomain of RHD1 are necessary for the interaction with RMB1 (Figures S3C and S3E). In the Y2H assay, RHD1 also interacts with itself, and this self-interaction is mediated through the zinc finger domain (Figures S3C and S3D).

### RMB1 binds to methylated DNA through the MBD domain

*Arabidopsis* has 13 canonical MBD proteins (Zemach and Grafi 2007). Our analysis indicates that *Arabidopsis* also has three non-canonical MBD proteins, namely RMB1, IDM1, and the protein encoded by AT4G14920 (Figure S4A). The IDM complex contains two MBD domain proteins, i.e., the histone acetyltransferase IDM1 and MBD7 (Qian et al. 2012; Lang et al. 2015). The two MBD domain proteins recognize methylated cytosine and thus ensure that the IDM histone acetyltransferase complex is directed only to methylated sequences (Qian et al. 2012; Lang et al. 2015). Many of the amino acid residues in the MBD domain of RMB1 are not conserved compared to the other MBD proteins (Figure S4B). We carried out electrophoretic mobility shift assays (EMSA) to determine whether RMB1 can bind to methylated DNA. The EMSA assays showed that RMB1 is capable of binding to DNA probes containing CG, CHG, or CHH methylation, and that the binding was competitively blocked by unlabeled methylated DNA but to a lower extent by unmethylated DNA of the same sequence (Figures 4A and 4B). To confirm RMB1 binding to methylated DNA, we carried out microscale thermophoresis (MST) assays and found that the MBD domain of RMB1 is capable of binding to DNA methylated in any sequence context (Figures 4C). We constructed four mutant versions of the MBD domain, including W22G, Y38F, T49A, and K50T (Figure S4B), and found that the W22G, Y38F, and T49A mutations decreased the methyl-DNA binding activity, while the K50T mutation abolished the methyl-DNA binding activity (Figures 4C). Expression of the wild-type but not of the K50T mutant version of RMB1 under its native promoter complemented the At5g39160 DNA hypermethylation phenotype of the *rmb1-1* mutant (Figure S5A). These results show that RMB1 is a novel methyl-DNA-binding protein, and that its methyl-DNA binding is critical for it to function in active DNA demethylation.

**Figure 4.**
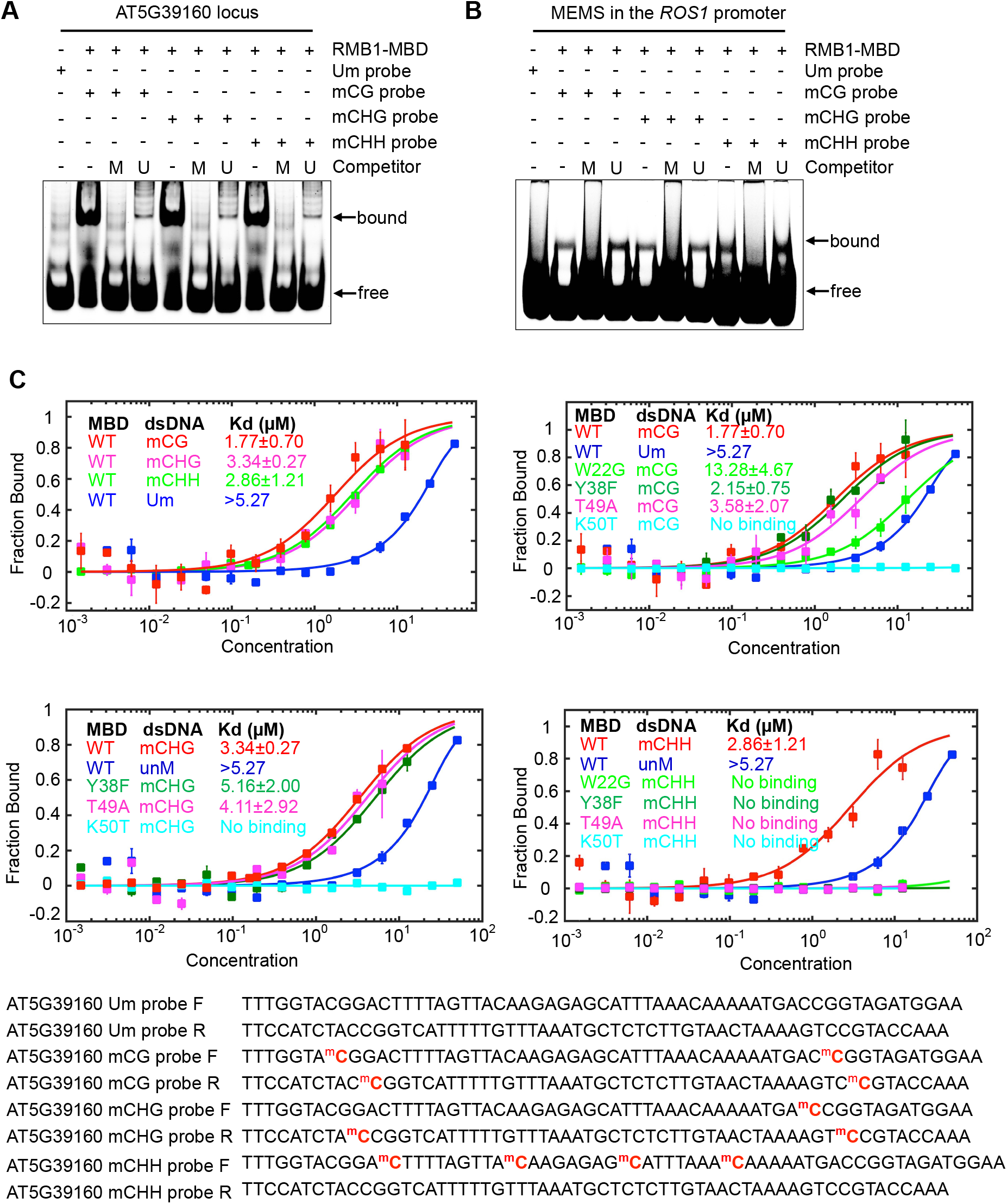
RMB1 binds to methylated DNA. **(A**,**B)** Electrophoretic mobility shift assay (EMSA) showing that the MBD domain of RMB1 (residues 1 to 81) binds to methylated oligonucleotides corresponding to the AT5G39160 locus (A) and the MEMS in the *ROS1* promoter (B). M, methylated; U, unmethylated. **(C)** RMB1 binding to labeled methyl-DNA sequence corresponding to the AT5G39160 locus assessed by microscale thermophoresis (MST). MST binding assays were used to quantify the interaction of RMB1 with an unmethylated (Um) probe or probes methylated (m) in CG, CHG, or CHH contexts. The W22G, Y38F, T49A, or K50T mutations in conserved residues in the MBD domain abolished or diminished the binding to methylated DNA. The detailed sequence information in each probe is displayed. The cytosine is labeled in the CHH context in the forward strand of the mCHH probe, but not in the complementary strand. Values are means ± SD (n=3). Curves indicate calculated fits; calculated binding affinities are displayed.

### RWD40, RMB1, and RHD1 regulate *ROS1* expression

To begin to explore the function of the protein complex formed by RWD40, RMB1, RHD1, and ROS1 in plants, we assessed the DNA methylation levels at the *ROS1* promoter region, which contains a DNA methylation monitoring sequence that senses methylation and demethylation activities and that regulates *ROS1* expression (Lei et al. 2015; Williams et al. 2015). Individual locus bisulfite sequencing showed that the DNA methylation level at the MEMS was increased in *rwd40-1, rmb1-1, rhd1-1*, and *ros1-4* compared to the wild-type control (Figure 5A). In contrast, the MEMS DNA methylation level was not increased in the *idm1-2* mutant (Figure 5A). Consistent with the changes in DNA methylation levels at the MEMS, *ROS1* expression was increased in *rwd40, rmb1*, and *rhd1* mutant plants but not in *idm1-2* mutant plants (Figure 5B). In the *ros1-4* mutant, *ROS1* expression measured using the 5’ primer pair (Figure 5B) was also increased, as expected (Lei et al.,2015). The expression of the *3xFLAG-RWD40* transgene driven by the *RWD40* native promoter in the *rwd40-1* mutant restored the expression of *ROS1* to the wild-type level (Figure 5B). ChIP-qPCR assays confirmed that ROS1, RWD40, RMB1, and RHD1 were enriched at the MEMS in the *ROS1* promoter (Figure 5C), and that *RWD40, RMB1*, and *RHD1* were necessary for ROS1 binding to the MEMS region (Figure 5D). Taken together, these results indicate that *RWD40, RMB1*, and *RHD1* mediate binding of ROS1 to the MEMS region, hence regulating its methylation level and the subsequent expression of *ROS1*.

**Figure 5.**
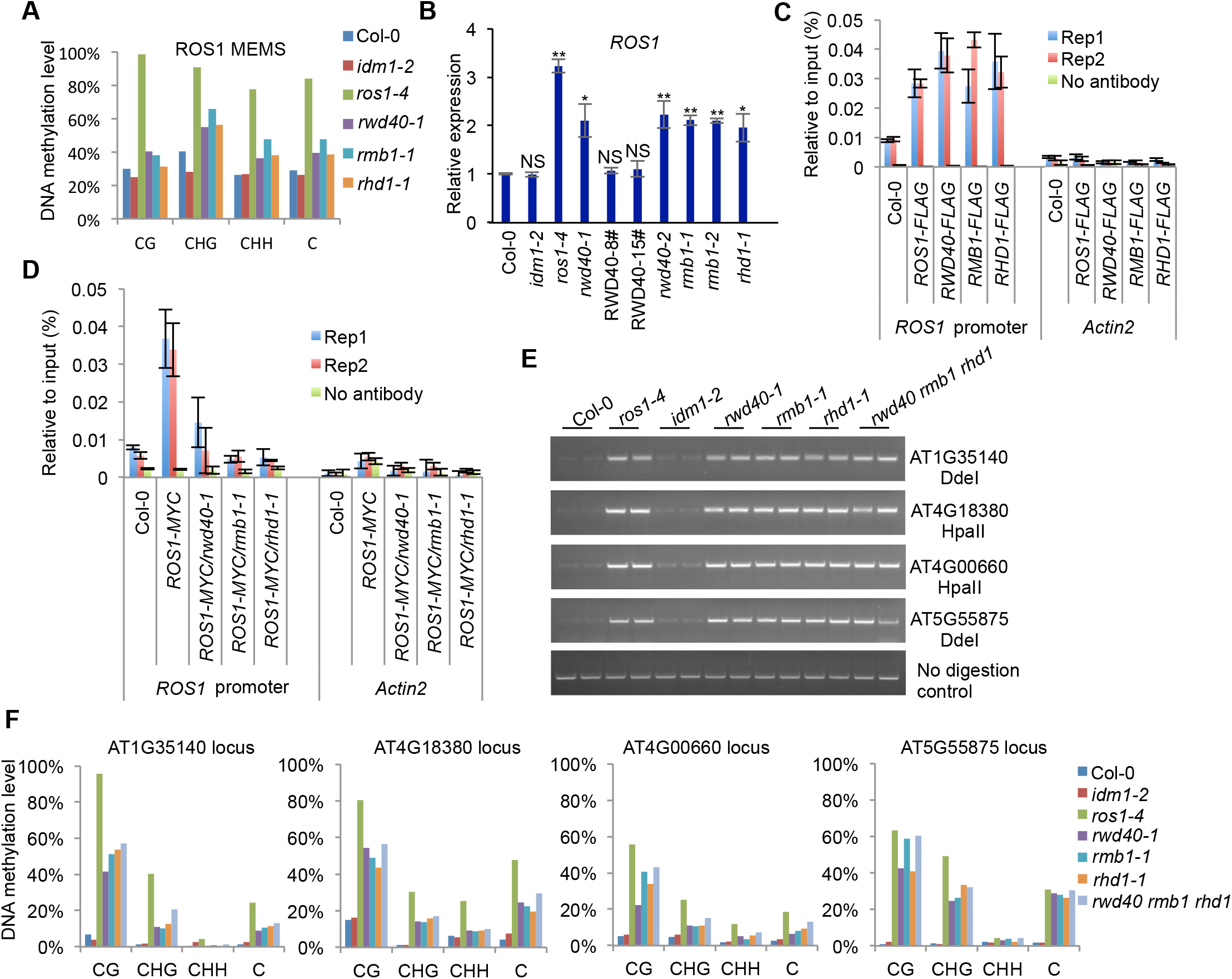
RWD40, RMB1, and RHD1 function in active DNA methylation. **(A)** Bisulfite sequencing data showing DNA methylation in different sequence contexts at the MEMS in the *ROS1* promoter in wild-type Col-0 and the indicated mutants. **(B)** Relative expression of *ROS1* in the indicated samples. Values are means ± SD of three biological replicates. *P < 0.05, **P < 0.01, compared with wild-type Col-0 plants; NS, not significantly different compared with wild-type Col-0 plants (2-tailed t test). **(C)** Enrichment of RWD40, RMB1, RHD1, and ROS1 at the MEMS in the *ROS1* promoter detected by ChIP-qPCR assays. The ChIP signal was quantified relative to input DNA. The no antibody precipitates served as negative control. *Actin 2* was used as a control region. Values are means ± SD of three technical replicates. Results of two biological replicates are displayed. “Rep” indicates replicate. **(D)** Effect of mutations in *RWD40, RMB1*, and *RHD1* on ROS1 protein enrichment at the MEMS in the *ROS1* promoter by ChIP-qPCR assays. **(E)** Chop-PCR showing that *rwd40-1, rmb1-1, rhd1-1*, and *ros1-4* mutants displayed an increased methylation phenotype at the indicated loci. Amplification of non-digested DNA served as a control. **(F)** Analysis of DNA methylation levels at several DNA demethylation target loci in wild-type Col-0 control plants and the indicated mutants by individual locus bisulfite sequencing analysis.

### RWD40, RMB1, and RHD1 regulate DNA demethylation in an IDM1-independent manner

*RWD40, RMB1, RHD1*, and *ROS1*, but not *IDM1*, are necessary for the prevention of DNA hypermethylation at the MEMS in the *ROS1* promoter (Figure 5A). These results suggest that RWD40, RMB1, and RHD1 might regulate ROS1-mediated DNA demethylation in a manner independent of IDM1. To further investigate the role of RWD40, RMB1, and RHD1 in the regulation of active DNA demethylation, we used Chop-PCR to assess the DNA methylation levels in *rwd40-1, rmb1-1*, and *rhd1-1* single mutants and in the *rwd40 rmb1 rhd1* triple mutant, as well as in *idm1-2*, at several genomic targets of ROS1. Similar to DNA methylation levels in the *ros1-4* mutant, DNA methylation levels at the AT1G35140, AT4G00660, AT4G18380, and AT5G55875 loci were increased in *rwd40-1, rmb1-1*, and *rhd1-1*, but not *idm1-2* mutant plants, relative to the Col-0 control (Figure 5E). Individual locus bisulfite sequencing data confirmed that the DNA methylation levels at these ROS1 targets were increased in *rwd40-1, rmb1-1, rhd1-1*, and *ros1-4* mutant plants, but not in *idm1-2* mutant plants (Figure 5F). Analysis of the *rwd40 rmb1 rhd1* triple mutant indicated that the *rwd40-1, rmb1-1*, and *rhd1-1* mutations are not additive in causing the DNA hypermethylation phenotype (Figures 5F). We also determined the DNA methylation levels at several genomic regions (including DT-75, DT-76, DT-77, and DT-78) that are known to be hypermethylated in *idm1* and *ros1* mutants (Qian et al. 2012)and that are amenable to Chop-PCR assays (Duan et al. 2017). DNA methylation levels were not increased at these regions in *rwd40-1, rmb1-1*, or *rhd1-1* mutant plants (Figure S6A). Together, these results support the inference that RWD40, RMB1, and RHD1 function in regulating active DNA demethylation independently of IDM1.

ROS1 and the IDM complex are required for the prevention of silencing of the *35S:SUC2* reporter gene in *Arabidopsis* (Qian et al. 2014; Lang et al. 2015; Duan et al. 2017; Nie et al. 2019). When we introgressed the *35S:SUC2* reporter gene into *rwd40-1, rmb1-1*, or *rhd1-1* mutant plants through genetic crosses, we found that the reporter gene was not silenced in any of these mutants (Figures S6B and S6C). Therefore, unlike IDM1 or ROS1, RWD40, RMB1, and RHD1 are not required for the prevention of silencing of the *35S:SUC2* reporter gene in *Arabidopsis*. We examined the deposition of H2A.Z and H3K18ac in wild-type *35S:SUC2* plants (Nie et al., 2019) at the 5 loci shown in Figure 5F, where DNA methylation levels were increased in RW40 complex-related mutants, but not in *idm1-2* (Figures 5E and 5F). Unlike the moderate co-existence of H2A.Z and H3K18ac in the IDM1-dependent AT1G62760 locus, only at the AT4G18380 locus were H2A.Z and H3K18ac both less deposited; at the AT5G55875 locus, H2A.Z and H3K18ac were rarely deposited; either H2A.Z or H3K18ac was deposited at the MEMS, AT1G35140, and AT4G00660 loci (Figure S6D). No consistent change in histone acetylation or H2A.Z deposition could be found at these 5 selected loci, which is in line with the notion that RWD40, RMB1, and RHD1 function in an IDM1-independent manner.

### The RWD40 complex prevents the silencing of several endogenous genes and regulates antibacterial resistance

To test the effect of the dysfunction of the RWD40 complex on genome-wide transcript levels, we profiled the transcriptomes of *ros1-4* and *rwd40 rmb1 rhd1* mutant plants by mRNA-seq. There were 3,817 and 2,970 up-regulated genes and 4,028 and 2,918 down-regulated genes in *rwd40 rmbd1 rhd1* triple-mutant and *ros1-4* mutant plants,respectively, compared to wild-type Col-0 plants (Figure 6A and Table S2). Of these, 1,673 up-regulated and 1,941 down-regulated genes were shared between *rwd40 rmbd1 rhd1* and *ros1-4* (Figure 6B). Several commonly down-regulated DEGs in *rwd40 rmbd1 rhd1* and *ros1-4* mutants were confirmed by qRT-PCR (Figure 6C). These results suggest that the RWD40 complex is required for the expression of some endogenous genes.

**Figure 6.**
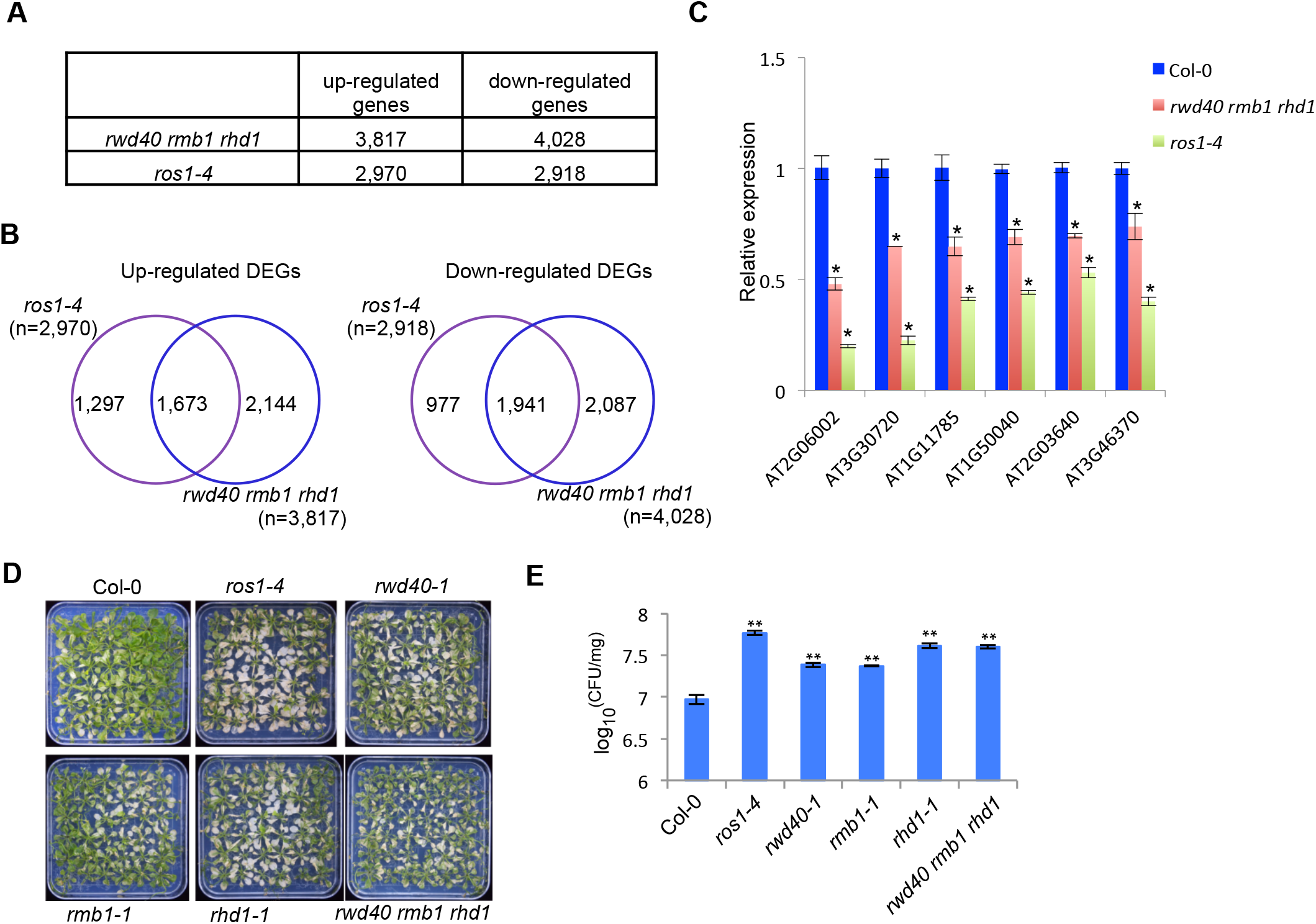
The RWD40 complex regulates gene expression and antibacterial resistance. **(A)** Number of differentially expressed genes identified in *rwd40 rmb1 rhd1* and *ros1-4* mutants by mRNA-seq. **(B)** Overlap of up-regulated or down-regulated genes between *rwd40 rmb1 rhd 1* and *ros1-4* mutants. **(C)** Validation of the down-regulation of six genes in *rwd40 rmb1 rhd1* and *ros1-4* mutants by qRT-PCR. Values are means ± SD of 3 biological replicates relative to transcript levels in wild-type Col-0 plants. *P < 0.05, compared with Col-0 samples (2-tailed t test). **(D)** Phenotypes of Col-0, *ros1-4, rwd40-1, rmb1-1, rhd1-1*, and *rwd40 rmb1 rhd* plants inoculated with Pto DC3000. Pictures were taken at 3-d post-inoculation (dpi). **(E)** Bacterial counts in Col-0 and the indicated mutants plants at 3 dpi. **P < 0.01, compared with Col-0 samples (2-tailed t test).

Exposure to the bacterial pathogen *Pseudomonas syringae* pv. *tomato* DC3000 (*Pto* DC3000) causes moderate but widespread differential DNA methylation in *Arabidopsis* (Dowen et al. 2012). A mildly enhanced bacterial growth was observed in *ros1* plants treated with the bacterial pathogen *Pto* DC3000, supporting a role of ROS1-dependent DNA demethylation in antibacterial resistance (Yu et al. 2013). To test whether the RWD40 complex regulates antibacterial resistance, we first inoculated the RWD40 complex mutants with *Pto* DC3000 by flood-inoculation assays (Ishiga et al. 2011). We found that similar to *ros1-4* mutant plants, *rwd40-1, rmb1-1*, and *rhd1-1* mutant plants were more susceptible to *Pto* DC3000 than wild-type plants (Figures 6D and 6E). These results point at a biological relevance of the RWD40 DNA demethylation complex in *Arabidopsis* antibacterial defense.

## DISCUSSION

In this study, we identified RWD40, RMB1, and RHD1 as cellular factors critical for the regulation of *ROS1* expression and for the prevention of DNA hypermethylation at several endogenous genomic regions (Figure 5). DNA methylation patterns are important for organismal development, carcinogenesis, and many other diseases, as well as for human aging (Jung and Pfeifer 2015; Klutstein et al. 2016). It is therefore important to understand how DNA methylation patterns are controlled by active DNA demethylation (Penterman et al. 2007; Furner and Matzke 2011; Zhang et al. 2018). Although the biochemistry of ROS1-mediated enzymatic removal of DNA methylation has been extensively studied, the mechanisms by which the enzymatic machinery is regulated and recruited to specific target sites are poorly understood (Qian et al. 2012; Lang et al. 2015; Nie et al. 2019). Our results indicate that RWD40, RMB1, RHD1, and ROS1 form a RWD40 complex, and that this protein complex contributes to locus-specific DNA demethylation. Moreover, ROS1 binding to the *ROS1* promoter requires RWD40, RMB1, and RHD1, suggesting that the RWD40 complex directs ROS1 targeting to this locus to regulate the expression of *ROS1*. Thus, the activity of the RWD40 complex at target genomic loci is likely self-restrained by *ROS1* expression level through its demethylation function at the MEMS site on the *ROS1* gene promoter (Figure 7). These findings provide novel insights into the regulation and targeting of active DNA demethylation in plants. In mammals, active DNA demethylation is initiated by the Tet dioxygenases (Wu and Zhang 2017). Several transcription factors have been shown to interact with and to target Tet enzymes to genes critical for cell differentiation and reprogramming (Costa et al. 2013; de la Rica et al. 2013; Wang et al. 2015; Xiong et al. 2016; Sardina et al. 2018). In *Arabidopsis*, the IDM histone acetyltransferase complex is required for directing ROS1 to a subset of active DNA demethylation target regions (Qian et al. 2012; Qian et al. 2014; Lang et al. 2015; Duan et al. 2017; Nie et al. 2019). MBD7 in the IDM complex recognizes dense mCpG sites and thus ensures that ROS1 is eventually targeted to genomic regions with densely methylated CpG sequences (Lang et al. 2015). On the other hand, the SANT/Myb/trihelix DNA-binding motif-containing protein HDP2 in the IDM complex probably helps direct the complex to possible regulatory sequences, thus avoiding heavily methylated transposon body regions and some genic regions that should not be demethylated (Duan et al. 2017). The IDM complex creates histone acetylation marks that attract the SWR complex, which then deposits the histone variant H2A.Z. Once deposited at the targeted site, H2A.Z helps to recruit ROS1 through a direct physical interaction (Nie et al. 2019). The protein complex identified in this study is similar to the IDM complex in that it also contains a protein (i.e., RMB1) that recognizes methylated DNA, and a transcription factor-like protein (i.e., RHD1). RMB1 ensures that the complex is directed to methylated genomic regions, while RHD1 probably helps to target the complex to regulatory sequences. Like the IDM complex, the RWD40 protein complex is likely also important for targeting active DNA demethylation to specific genomic regions (Figures 5A, 5E, and 5F). Our results suggest that the RWD40 complex targets active DNA demethylation to genomic regions that are different from those targeted by the IDM complex. This difference can be explained by the fact that RMB1 recognizes mC marks in all sequence contexts (Figure 4), whereas MBD7 and IDM1 bind to mCpG sequences only (Qian et al. 2012; Lang et al. 2015) (Figure S5B and S5C). In addition, HDP2 and RHD1 likely have different sequence specificities in their interactions with DNA, which probably also contributes to the different targeting preferences of the two protein complexes. The lack of DNA hypermethylation at the AT5G39160 locus in *rhd1* mutant plants (Figure S2F) indicates that RHD1 may be dispensable for some targets of the RWD40 protein complex, either due to genetic redundancy with another transcription factor-like protein, or because its function is genomic region-dependent. It should be noted that H2A.Z deposition and histone acetylation are not restricted to the IDM1-dependent ROS1-targets, since H2A.Z or H3K18ac are also deposited at some loci targeted by RWD40 complex (Figure S6D); this observation suggests that other IDM1-independent acetylation mechanisms are associated with ROS1 activity and possibly involved in H2A.Z deposition.

**Figure 7.**
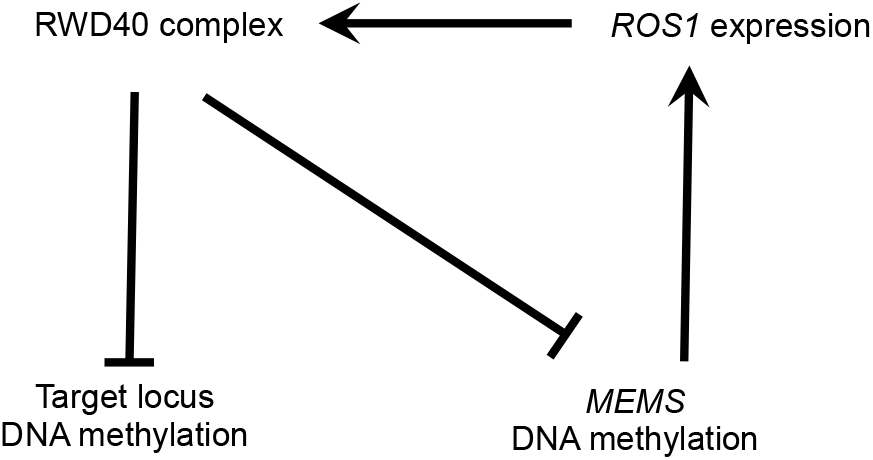
Schematic model showing that RWD40 may inhibit its own function by targeting the MEMS at the ROS1 promoter. The RWD40 complex decreases the DNA methylation level of the MEMS site at the *ROS1* promoter, causing reduced *ROS1* expression. The reduced *ROS1* expression would hinder the DNA demethylation function of the RWD40 complex at target genomic loci.

Another important difference between the RWD40 complex and the IDM complex is that the RWD40 complex contains ROS1 and thus has a direct role in targeting active DNA demethylation. Furthermore, because the RWD40 complex targets the MEMS in the *ROS1* promoter, it can regulate *ROS1* gene expression. We propose that through the regulation of *ROS1* gene expression, the RWD40 complex is expected to have a broad effect on genomic DNA methylation patterns. The genome-wide effects of DNA demethylation factors can be difficult to analyze because DNA methylation changes caused by mutations in these factors are often quite subtle (Nie et al. 2019). Future studies are needed to determine the genome-wide effects of *rwd40, rmb1*, and *rhd1* mutations, and to identify the direct targets of the RWD40 complex at the whole genome level.

The data presented here, together with our previous work (Lang et al. 2015; Nie et al. 2019), suggest that active DNA demethylation activities are targeted only to methylated genomic regions by proteins that bind to methyl-DNA. Such targeting would be biologically relevant, because only methylated genomic regions may require demethylation. Our findings also suggest that the presence of a transcription factor-like protein is important for the targeting of active DNA demethylation. It is tempting to speculate that these transcription factor-like proteins preferentially bind to regulatory sequences, and thus help target active DNA demethylation to regulatory sequences in order to prevent harmful silencing and precisely regulate gene expression.

## MATERIALS AND METHODS

### Plant materials and growth conditions

Plants were grown under long-day conditions (16-h light/8-h dark) at 22°C. The T-DNA insertion mutants *rwd40-1* (SALK_068825), *rwd40-2* (SALK_012947), *rmb1-1* (SALK_110885), *rmb1-2* (SALK_100783), *rhd1-1* (SALK_092897), *ros1-4* (SALK_045303), and *idm1-2* (SALK_062999) were obtained from the *Arabidopsis* Biological Resource Center (http://www.arabidopsis.org) and genotyped by PCR with the primers listed in Table S3.

## METHODS DETAILS

### Mutant plant complementation

For the complementation of mutants, the *RWD40, RMB1*, or *RHD1* genomic DNA with the 2-kb upstream region (as the native promoter region) was amplified from genomic DNA of Col-0 with the primers listed in Table S3. The amplification products were cloned into the pCambia1305 vector (with a *3xFLAG* tag or a *3xMYC* tag at the C-terminus) by T4 DNA ligase (NEB). The constructs were transformed into mutants using *Agrobacterium tumefaciens* GV3101 by the standard floral dip method (Clough and Bent 1998). Unsegregated T3 homozygous complementation lines were then used for further experiments.

### Real-time qPCR and Chop-PCR

For real-time qPCR analysis, total RNA was extracted from 0.1 g of 14-day-old *Arabidopsis* seedlings with the RNeasy plant kit (Qiagen). 2 μg of the mRNA were converted into cDNA with the First-Strand cDNA Synthesis Kit (TRNSGENE Bio., China) following the manufacturer’s instructions, and the cDNAs were used as templates for real-time PCR with iQ SYBR green supermix (Bio-RAD). For Chop-PCR, genomic DNA was extracted from 14-day-old seedlings, and 500 ng of DNA were digested with the indicated methylation-sensitive enzyme in a 20-μL reaction mixture. After digestion, PCR was performed using 2 μl of the digested DNA as template in a 20-μl reaction mixture with the primers listed in Table S3. Undigested DNA was amplified as the loading control.

### IP and LC-MS/MS analysis

For immunoprecipitation (IP), about 5 g of floral tissue for each epitope-tagged transgenic line were used. Dynabeads (10004D, Invitrogen) conjugated with FLAG antibody (F1804, Sigma) were applied for IP. Affinity purification was performed as previously described (Law et al. 2010), and the protein samples were subjected to Liquid Chromatography-Mass Spectrometry (LC-MS) /MS analysis as previously described (Lang et al. 2015; Qin et al. 2017; Nie et al. 2019).

### Protein extraction and western blot detection

About 0.1 g of plant tissue were harvested and ground to a fine power in liquid N_2_. Total protein was extracted by protein lysis buffer (LB: 0.5 mM DTT, 5 mM MgCl_2_, 50 mM Tris (pH 7.6), 10% glycerol, 150 mM NaCl, 0.1% NP-40, 1 mM PMSF, and protease inhibitor cocktail (Roche)). The proteins were then detected by western blot with anti-FLAG (F1804, Sigma) or anti-MYC (05-724, Millipore).

### Split luciferase complementation assays

The coding sequences of RWD40, RMB1, RHD1, and ROS1 proteins were cloned into pCAMBIA-cLUC and/or pCAMBIA-nLUC vectors. *A. tumefaciens* GV3101 carrying different constructs were infiltrated into 4-week-old *N. benthamiana* leaves. Luciferase activity was detected 48 h post infiltration.

### Individual locus bisulfite sequencing

200 ng of genomic DNA extracted from Col-0 wild type and the indicated mutants were treated with the BisulFlash DNA Modification Kit (Epigentek) according to the manufacturer’s protocol. A 2-μL volume of bisulfite-treated DNA was used for PCR amplification with the primers listed in Table S3. The PCR products were cloned into the pMD18-T vector (Takara) according to the supplier’s instructions. At least 15 independent clones were sequenced per sample and analyzed using the online tool Kismeth (http://katahdin.mssm.edu/kismeth/revpage.pl).

### Bimolecular fluorescence complementation (BiFC) assay

The coding sequences of RWD40, RMB1, RHD1, and ROS1 were cloned into the p2YN or p2YC vector to generate fused split YFP constructs. For protein-protein interaction analysis, *A. tumefaciens* GV3101 carrying the indicated constructs were cultured overnight. When the cultures reached an OD_600_ of 1.0, they were firstly collected by centrifuge and then were resuspended in buffer containing 10 mM MgCl_2_, 100 µM acetosyringone, and 10 mM MES (pH 5.6). The suspensions were kept at room temperature for 2 hours and then were infiltrated into *N. benthamiana* leaves. Fluorescence was examined at 2 days post infiltration.

### Gel filtration assays and western blot analysis

Western blotting of gel filtration samples was performed as previously described (Duan et al. 2017). In brief, 2 g of flower tissue from Col-0 wild type or from the indicated transgenic plants were harvested and ground into a fine power in liquid N_2_. The fine powder was resuspended in 3 ml of lysis buffer (0.5 mM DTT, 5 mM MgCl_2_, 50 mM Tris [pH 7.6], 10% glycerol, 150 mM NaCl, 0.1% NP-40, 1 mM PMSF), and the suspension was kept at 4°C for 5 min without shaking. The supernatant was loaded into a HiPrep 16/60 Sephacryl S-300 HR column (GE Healthcare), and 10 fractions were collected. Each fraction was run on 10% SDS-PAGE for western blot detection.

### Y2H assay

In brief, the corresponding cDNA sequences were cloned into pGADT7-AD or pGBKT7-BD vectors (Clontech), and the pair of genes to be tested for interaction were co-transformed into the yeast strain Gold (Clontech). Y2H assays were performed as previously described (Bai et al. 2013).

### Protein expression and purification

Arabidopsis *RWD40, RMB1*, and *RHD1* were cloned into the pFastBac1 vector adding a FLAG tag at their N-terminus. Arabidopsis *ROS1* (encoding amino acids 1-100) was cloned into the pFastBac1 vector adding a double His tag (6xHis-13aa-10xHis) fused with GFP at its N-terminus. Recombinant baculoviruses were generated by the Bac-to-Bac system in Sf9 insect cells. For protein co-expression, the insect cells were grown to a density of 2.0E+06 cells per mL and were then infected with four separate viruses. The infected cells were cultured for 60-72 h at 27°C before collection. Collected cell pellets were stored at -80°C before use. Each cell pellet was resuspended in buffer A (20 mM HEPES pH 7.5, 150 mM NaCl, 10% v/v glycerol, 50 mM imidazole, and 0.1 mM Tris (2-carboxyethyl) phosphine hydrochloride (TCEP), supplemented with protease inhibitor cocktail) and then lysed by sonication. The suspension was centrifuged at 18,000 × *g* for 30 min at 4°C. The supernatant was incubated with Ni-NTA resin at 4°C for 1 h. The resin was washed with 20 column volumes of buffer A in a gravity column. The complex was eluted by buffer B (buffer A with 250 mM imidazole) and was then loaded onto a Superdex200 10/300 GL column equilibrated with buffer containing 20 mM HEPES pH 7.5, 100 mM NaCl, and 0.1 mM TCEP. Fractions containing the ROS1 complex were collected and analyzed with SDS-PAGE and Coomassie brilliant blue staining.

### Microscale thermophoresis (MST)

The sequence encoding the MBD domain of RMB1 (amino acids 1-81) was cloned into a pET-Sumo vector to fuse the target protein with a hexahistidine plus yeast sumo tag at the N-terminus. The His-Sumo tagged MBD domain of RMB1 recombinant protein was transformed into *Escherichia coli* strain BL21 (DE3). When the OD_600_ of the cell culture reached 0.7, IPTG was added to a final concentration of 0.2 mM to induce protein expression. The cells were harvested by centrifugation, re-suspended in buffer A (500 mM NaCl, 20 mM Tris-HCl, pH 7.5), and then lysed by sonication. The supernatant was loaded into a HisTrap column (GE Healthcare) and washed with buffer A and buffer B (buffer A plus 50 mM immidazol). The target protein was finally eluted with buffer C (buffer A plus 500 mM immidazol). The eluted protein was dialyzed against buffer A and digested with Ulp1 protease. The His-Sumo tag was removed by passing the digestion product through a second step HisTrp column. The flow-through protein was dialyzed against buffer D (100 mM NaCl, 20 mM HEPES, pH 7.0, 5 mM DTT) and loaded onto a Heparin column (GE Healthcare). A linear gradient from buffer D to buffer E (2 M NaCl, 20 mM HEPES, pH 7.0) was applied to elute the target protein. The eluted protein was further purified using a Superdex G200 column with buffer E (50 mM NaCl, 20 mM HEPES, pH 7.0, 5 mM DTT) and concentrated to 10 mg/ml for further use. The mutated forms of the MBD domain of RMB1 were expressed and purified following the same protocol.

The MST assay was performed using a Monolith NT.115 instrument (NanoTemper Technologies) (Jerabek-Willemsen et al. 2014; Harris et al. 2018). The purified MBD domain of RMB1 was half-and-half diluted in 14 steps with the buffer E, covering a concentration range from about 50 μM to 1.5 nM (Figure 4C). For each DNA duplex, one DNA strand is labeled with Biotin at the 5’ during the synthesis (Figure 4C). The Biotin labeled strand and the unlabeled complementary strand were annealed together and further mixed with the diluted protein steps. The mixture was loaded into the MST capillaries. Three repeats for each measurement were conducted. The data were analyzed using the NanoTemper analysis software (NanoTemper Technologies).

### *Pto* DC3000 bacterial infection assays

Pto DC3000 bacterial infection assays were performed according to (Ishiga et al. 2011) with some modifications. Briefly, the indicated mutants were germinated and grown on 1/2 MS medium. A single colony of *Pto* DC3000 was inoculated onto new mannitol-glutamate medium agar with rifampicin (50 μg/ml) and streaked on the plate. The plate was inoculated at 28°C for 36 hours. The collected bacterial colonies were suspended in an appropriate volume of sterile distilled water. 40 ml of a *Pto* DC3000 bacterial suspension (4 × 10^6^ CFU, colony forming units) were poured onto the plate containing 3-week-old *Arabidopsis* seedlings and incubated for 2 minutes at room temperature. The bacterial suspension was then removed by pipetting and the plates were sealed with 3M Micropore 2.5 cm surgical tape. The infected seedlings were incubated in long-day (16-h light / 8-h dark) conditions at 22 °C for three days. The development of symptoms was evaluated and photographed at 3 days-post-inoculation (dpi). Bacterial growth was evaluated according to (Ishiga et al. 2011).

### mRNA-seq analysis

For mRNA-seq analyses, total RNA was extracted from 2-week-old seedlings with the RNeasy plant kit (Qiagen), and subjected to RNA deep sequencing. Three biological replicates were performed per genotype. The libraries were constructed and sequenced at the Genomics Core Facility of the Shanghai Centre for Plant Stress Biology, Chinese Academy of Sciences, using an Illumina HiSeq2500. Low-quality sequences (q < 20) and adapters were trimmed using Trimmomatic and cutadpte (Bolger et al. 2014). Clean reads were mapped to the *Arabidopsis* reference genome (TAIR10) using STAR (Dobin et al. 2013). DESeq2 was used to detect differentially expressed genes according to adjusted p value (padj) threshold of <0.01 (Love et al. 2014).

## SUPPLEMENTAL INFORMATION

Supplemental Information includes Figures S1-S7 and Tables S1-S3, and can be found with this article online.

## AUTHOR CONTRIBUTIONS

J.-K.Z. and P.L. designed the study. P.L., W.-F.N., X.X., Y.W., Y.J., P.H., X.L., G.Q., H.H., J.D., and Z.L. performed the experiments. P.L., W.-F.N., and J.-K.Z. analyzed the data. W.-F.N., R.L.-D., and J.-K.Z. wrote the manuscript.

## ACKNOWLEDGEMENTS

This work was supported by the Chinese Academy of Sciences, National Nature Science Foundation of China (32002046), and Natural Science Foundation of Jiangsu Province (BK20200948).

## FIGURE LEGENDS

**Figure S1.**
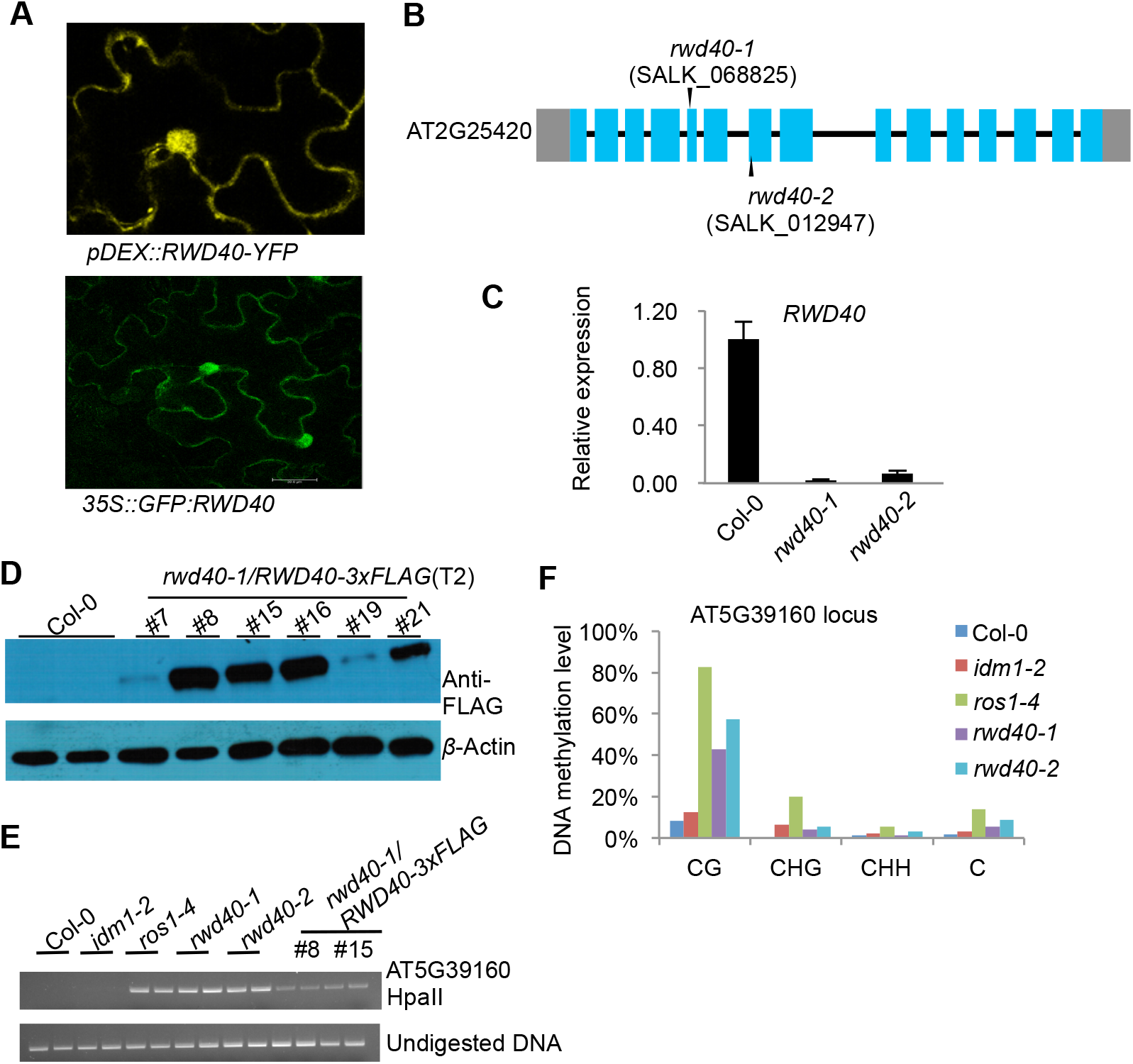
DNA demethylation phenotypes o*f rwd40-1* a*nd rwd40-2* mutants and mutant complementation. **(A)** RWD40 localization in *Nicotiana benthamiana*. **(B)** T-DNA insertion position in *rwd40-1* and *rwd40-2* mutants. Boxes and lines denote exons and introns, respectively. **(C)** Expression of *RWD40* in mutants relative to that in wild-type Col-0 control plants, as measured by qRT-PCR. Values are means ± SD of three biological replicates. **(D)** Western blot analysis of RWD40-3xFLAG in T2 *rwd40-1* transgenic lines. **(E)** DNA methylation levels in *rwd40* mutant alleles and complementation lines at the AT5G39160 locus as determined by Chop-PCR. **(F)** DNA methylation levels at the AT5G39160 locus in *rwd40-1* and *rwd40-2* mutants as determined by individual bisulfite sequencing.

**Figure S2.**
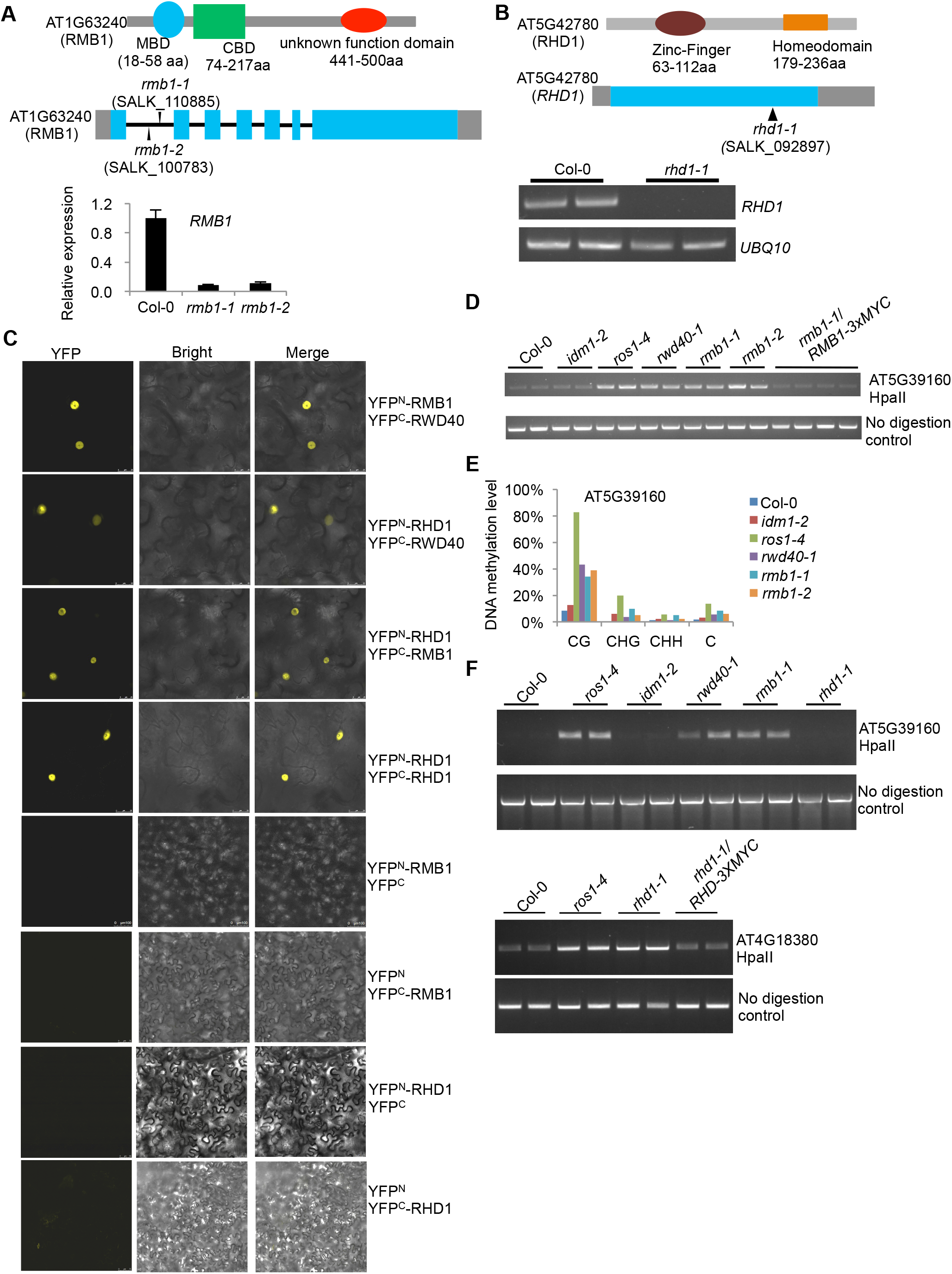
Characterization of RMB1 and RHD1. **(A)** RMB1 domains and mutants. Upper panel, schematic representation of the RMB1 protein. CBD, C (AT)-rich DNA-binding domain. Middle panel, T-DNA insertion positions in *rmb1-1* and *rmb1-2* mutants. Boxes and lines denote exons and introns, respectively. Lower panel, RMB1 transcript levels in the mutants. **(B)** Schematic representation of the RMB1 protein (upper panel), T-DNA insertion position in the *rhd1-1* mutant (middle panel), and RHD1 transcripts levels in the *rhd1-1* mutant by RT-PCR detection (lower panel). **(C)** Interactions among RWD40, RMB1, and RHD1 in BiFC assays. **(D)** DNA methylation levels in *rmb1* mutant alleles and in complementation lines at the AT5G39160 locus as determined by Chop-PCR. **(E)** DNA methylation levels at the AT5G39160 locus in *rmb1-1* and *rmb1-2* mutants as determined by individual locus bisulfite sequencing. **(F)** DNA methylation levels in wild-type Col-0 and indicated mutants at the AT5G39160 locus (upper panel), and in *rhd1* mutant alleles and complementation line at the AT4G18380 locus as determined by Chop-PCR (lower panel).

**Figure S3.**
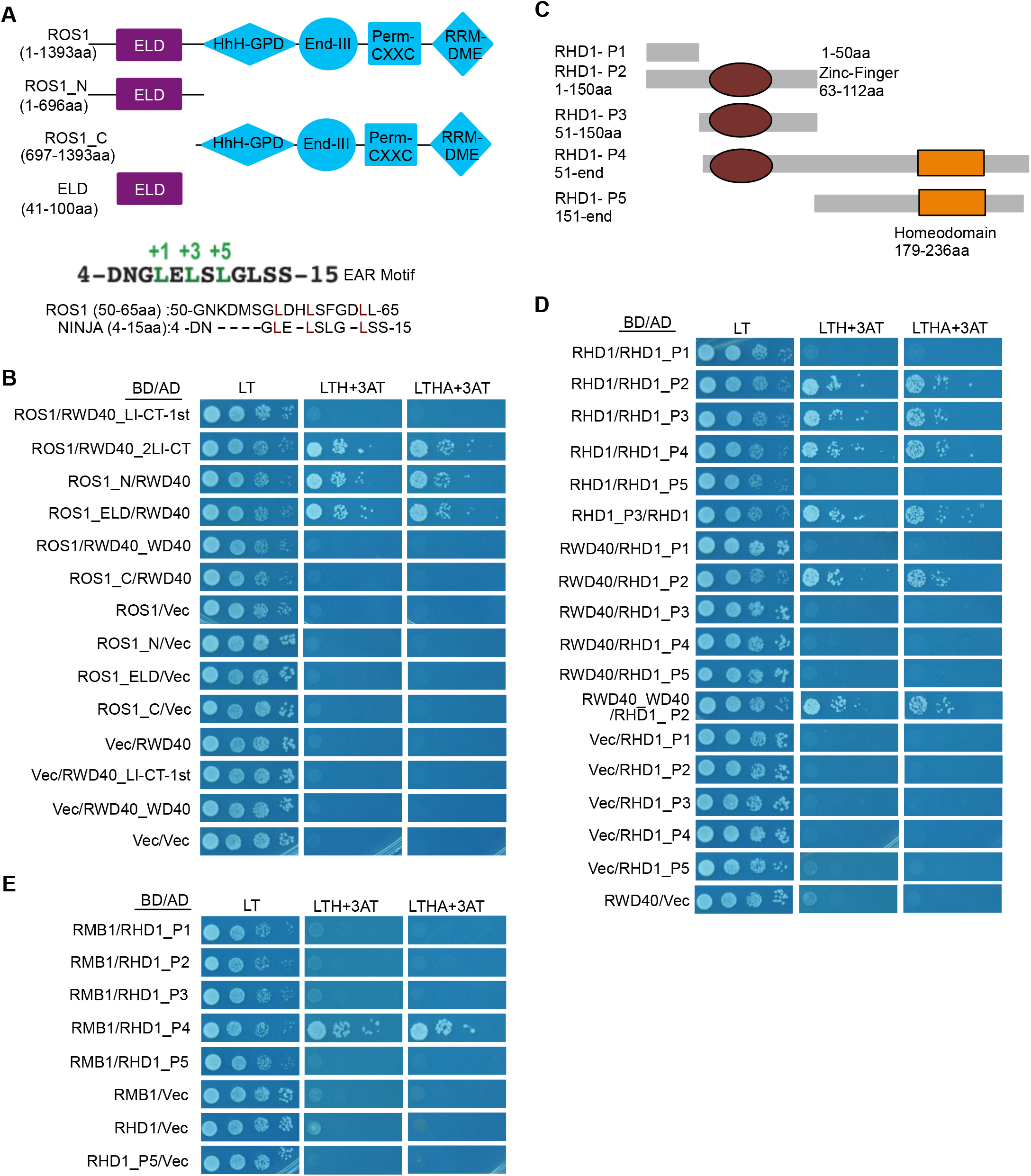
Interactions among RWD40, RMB1, RHD1, and ROS1 in Y2H assays. **(A)** Schematic representation of the truncated forms of ROS1 used in the Y2H assays and of the ELM domain identified in N-terminal regions of ROS1. HhH-GPD, hallmark Helix-hairpin-helix and Gly/Pro-rich loop domain; EndIIII, endonuclease III; Perm-CXXC, permuted version of a single unit of the zinc finger-CXXC; RRM_DME, RRM-fold domain present at the C-terminus of Demeter-like glycoslyases; EAR, ethylene response factor-associated amphiphilic repression domain; ELD, EAR-like motif. **(B)** The ELD domain of ROS1 interacts with RWD40 in Y2H assays. **(C)** Schematic representation of the truncated forms of RHD1 used in Y2H assays. **(D)** RHD1 interacts with RHD1 through the zinc-finger domain, and the WD40 domain of RWD40 interacts with the zinc-finger domain of RHD1 in Y2H assays. **(E)** RMB1 interacts with full-length RHD1 but does not interact with its zinc-finger domain or homeodomain.

**Figure S4.**
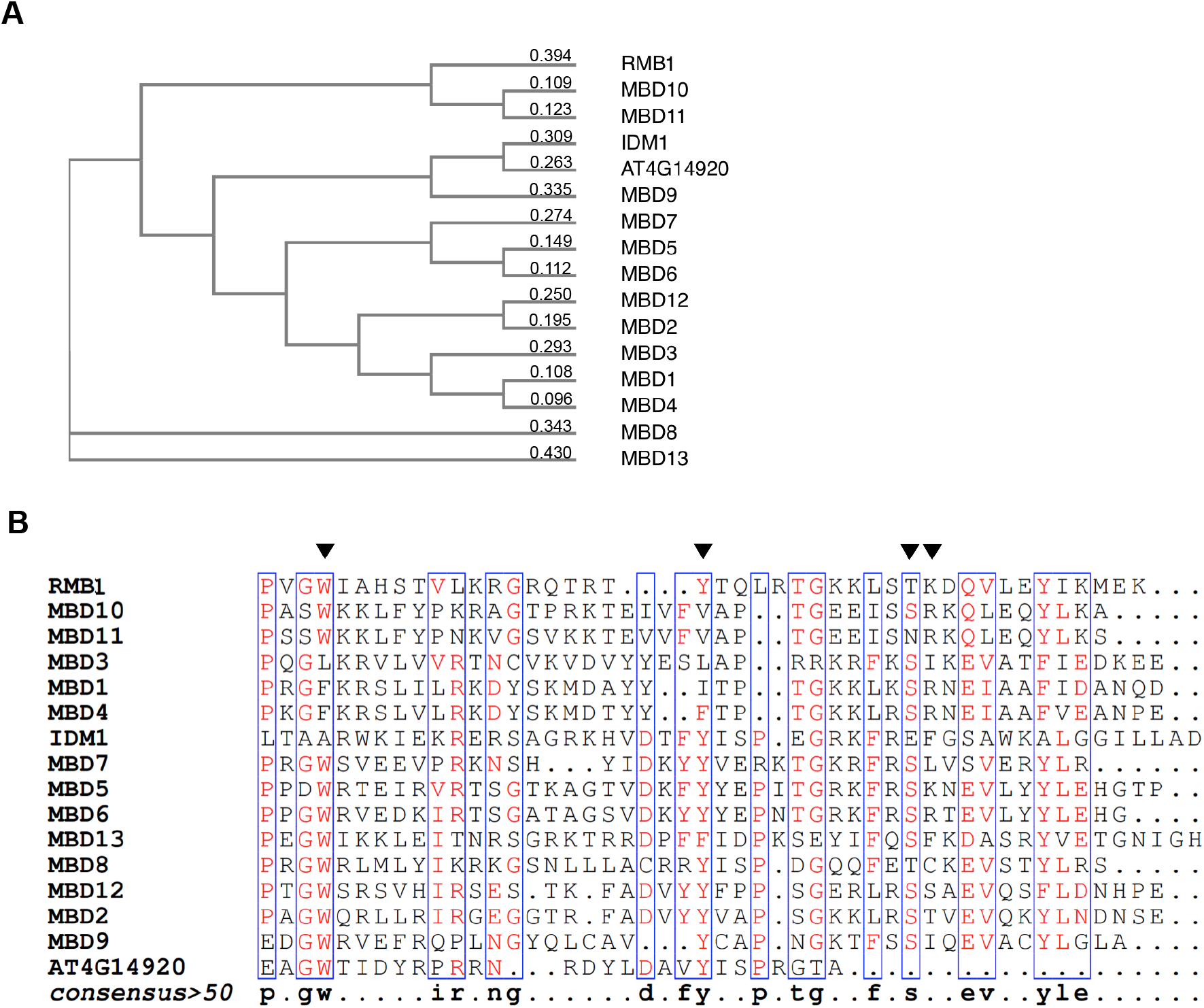
Analysis of MBD domain-containing proteins in *Arabidopsis*. **(A)** Phylogenetic tree of MBD domain-containing proteins in *A. thaliana*. **(B)** Alignment of MBD domains in MBD domain-containing proteins in *A. thaliana*. The amino acid residues targeted for mutations in the MBD domain are indicated by triangles.

**Figure S5.**
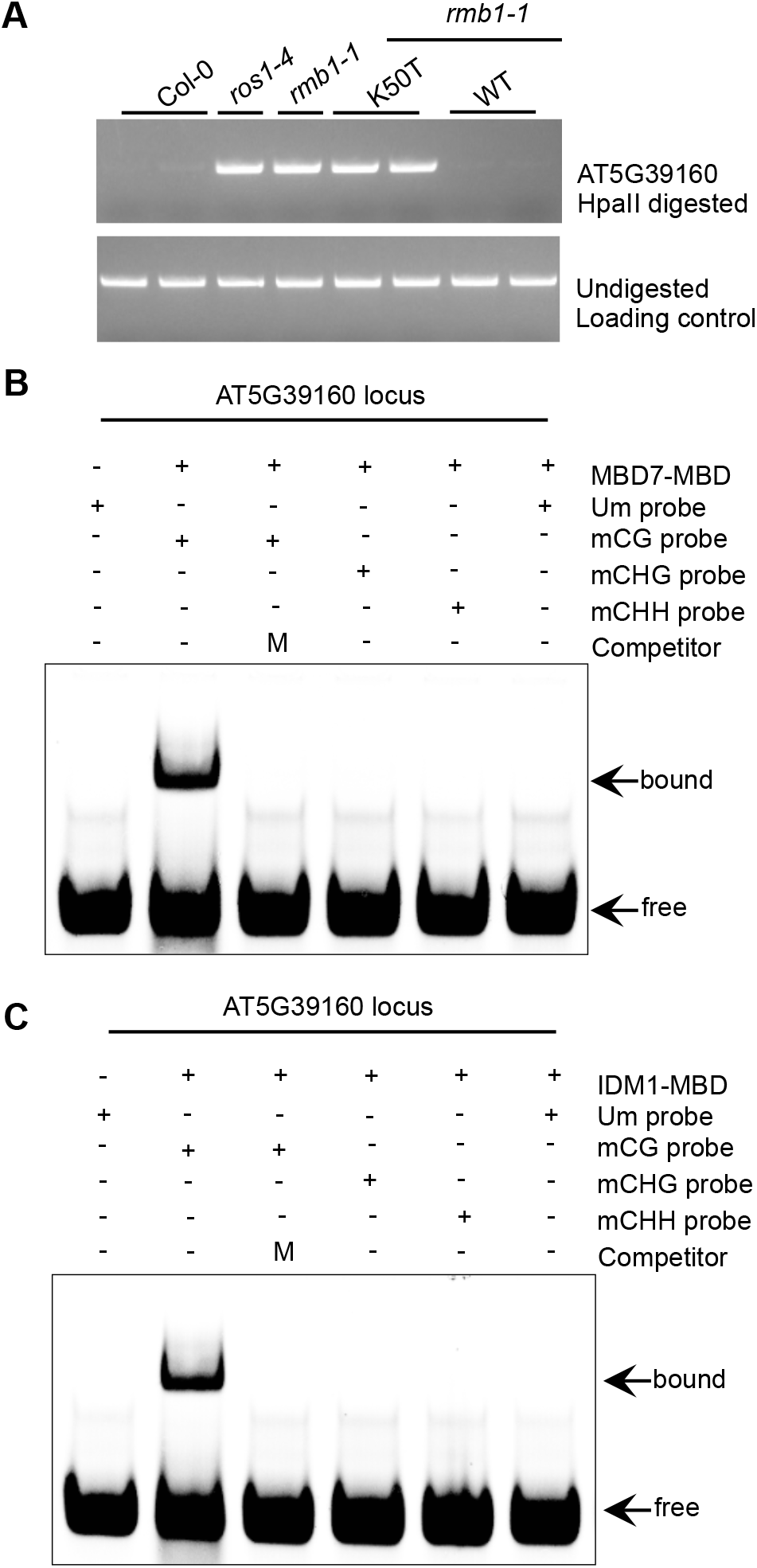
Mutation of the MBD domain reduces the binding of RMB1 to methylated DNA and abolishes the demethylation function of RMB1. **(A)** DNA methylation level at AT5G39160 in the *rmb1-1* mutant and in the *rmbd1-1* mutant expressing WT (wild-type) or K50T-mutated RMB1 as determined by Chop-PCR. **(B**,**C)** EMSA of the MBD domain of MBD7 (C) and IDM1 (D) binding to methylated oligonucleotides corresponding to the AT5G39160 locus. M, methylated. Um, unmethylated.

**Figure S6.**
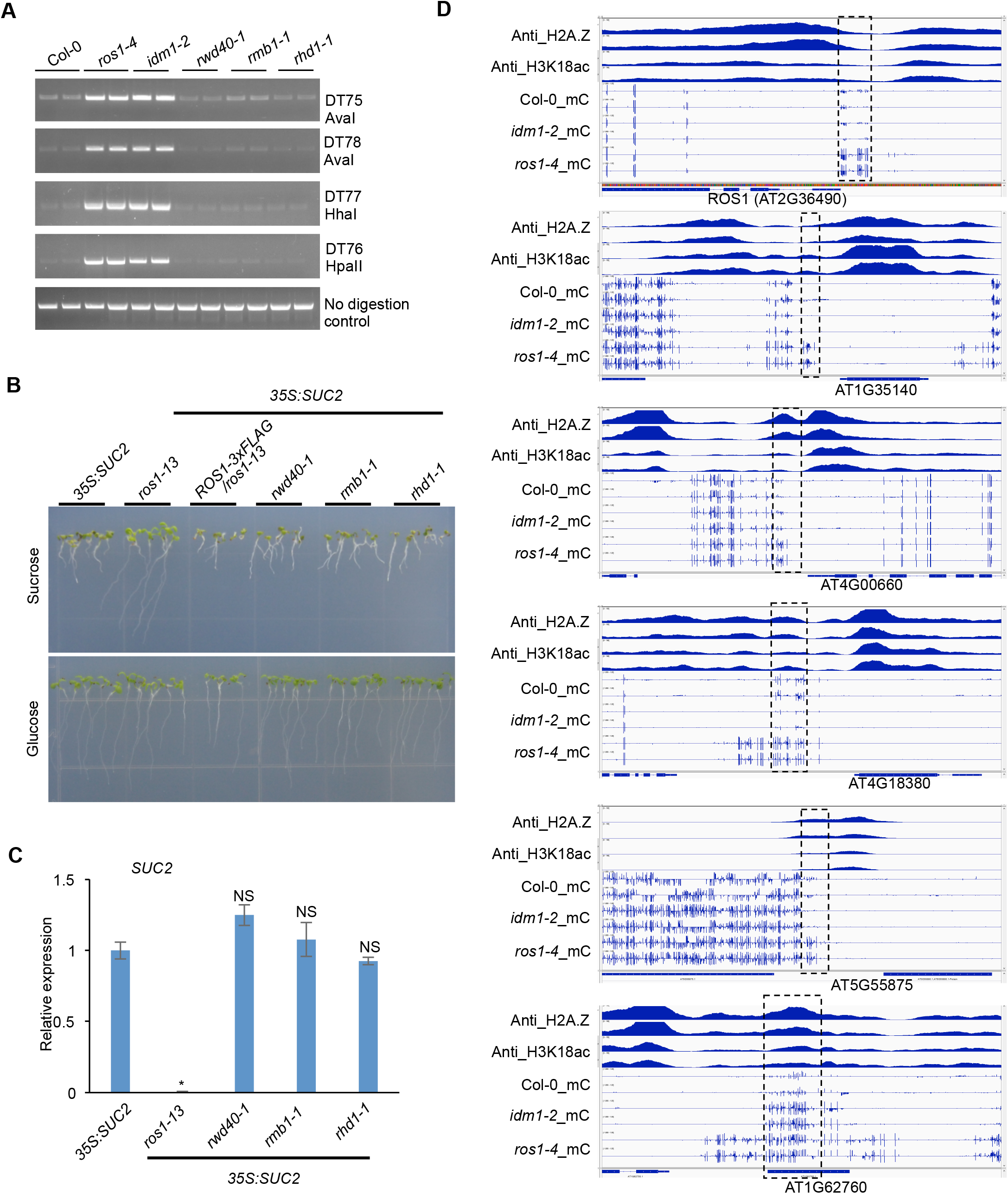
Dysfunction of RWD40, RMB1, or RHD1 does not cause silencing of the *35S:SUC2* transgene. **(A)** Chop-PCR showing that *rwd40-1, rmb1-1*, and *rhd1-1* mutants do not display an increased methylation phenotype in IDM1-dependent DNA demethylation target loci (Qian et al. 2012). Amplification of non-digested DNA served as a control. **(B)** Root phenotype in *rwd40-1, rmb1-1*, and *rhd1-1* in the *35S:SUC2* transgenic background. **(C)** Relative expression of the *35S:SUC2* transgene in the indicated mutants. Values are relative to transcript levels in *35S:SUC2* control plants and are means ± SD of three biological replicates. *P < 0.01, compared with *35S:SUC2* plants; NS, not significantly different compared with *35S:SUC2* plants (2-tailed t test). **(D)** Screenshots of DNA cytosine methylation status in Col-0, *idm1-2*, and *ros1-4*, and screenshots of H2A.Z deposition and H3K18Ac in wild-type control plants (Nie et al., 2019) at the tested loci in Figure 5. AT1G62760 is an *IDM1*-dependent active DNA demethylation target (Qian et al., 2012, Nie et al., 2019), and serves as a positive control of the co-existence of H2A.Z and H3K18ac.

**Figure S7.**
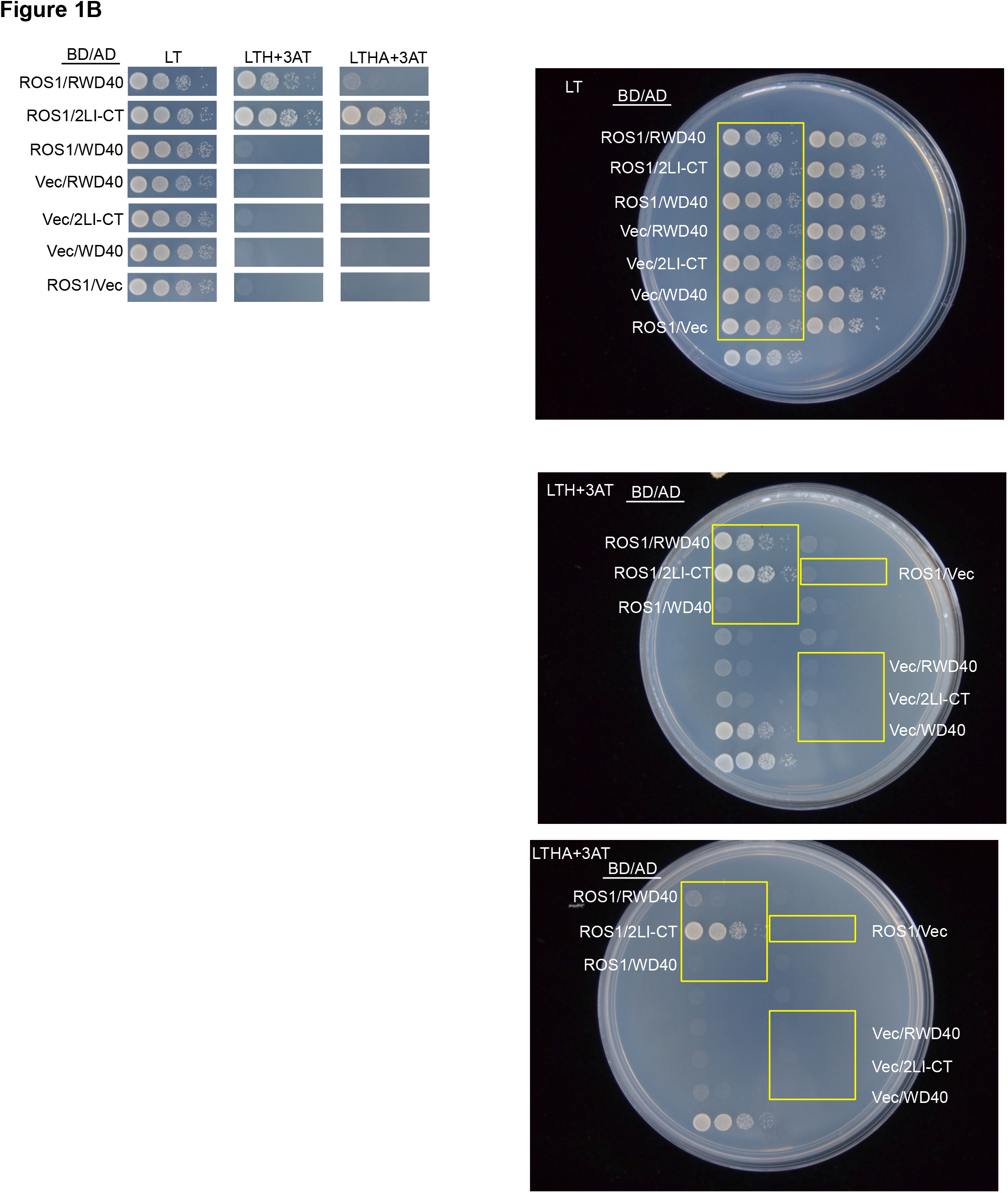
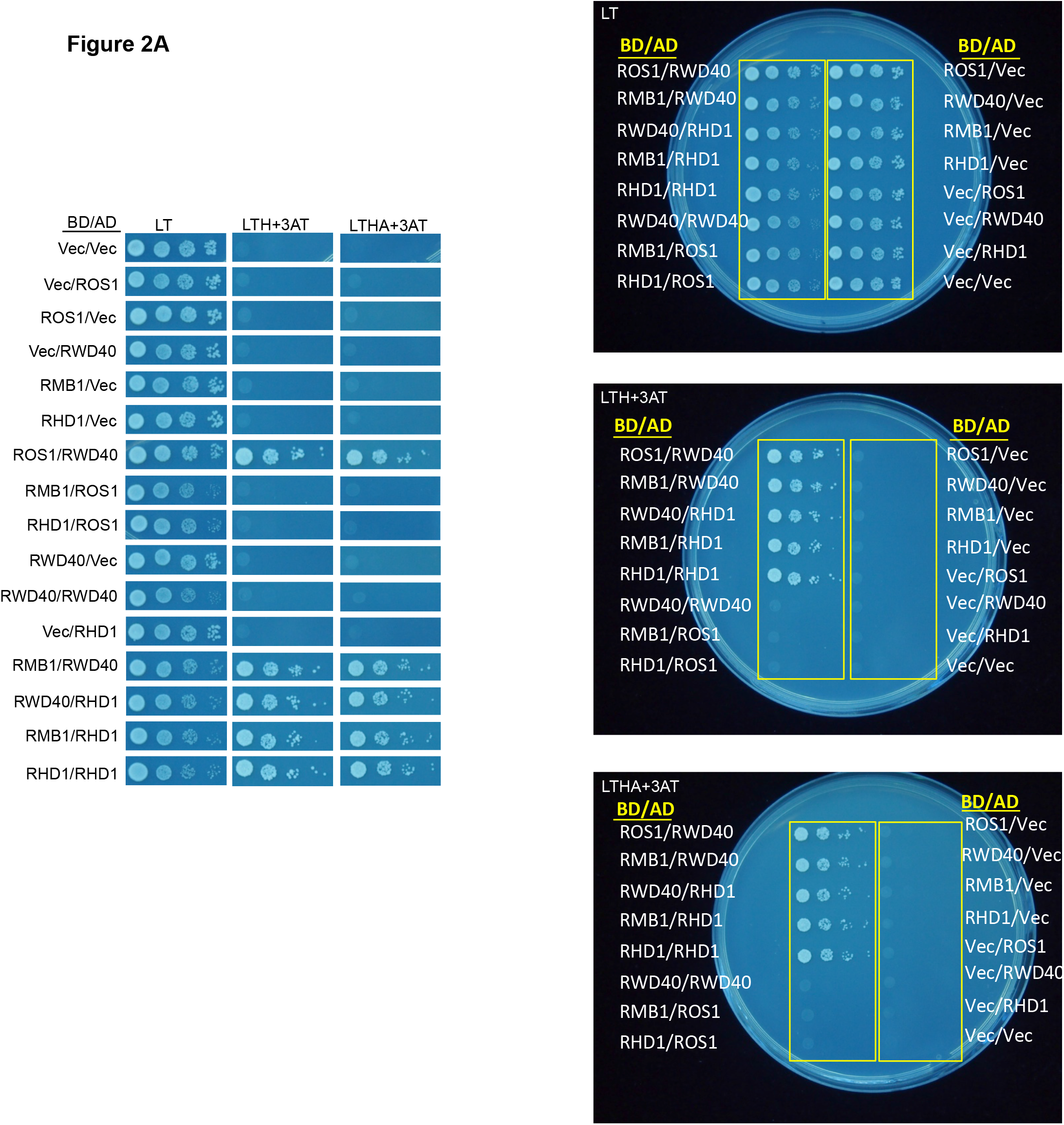
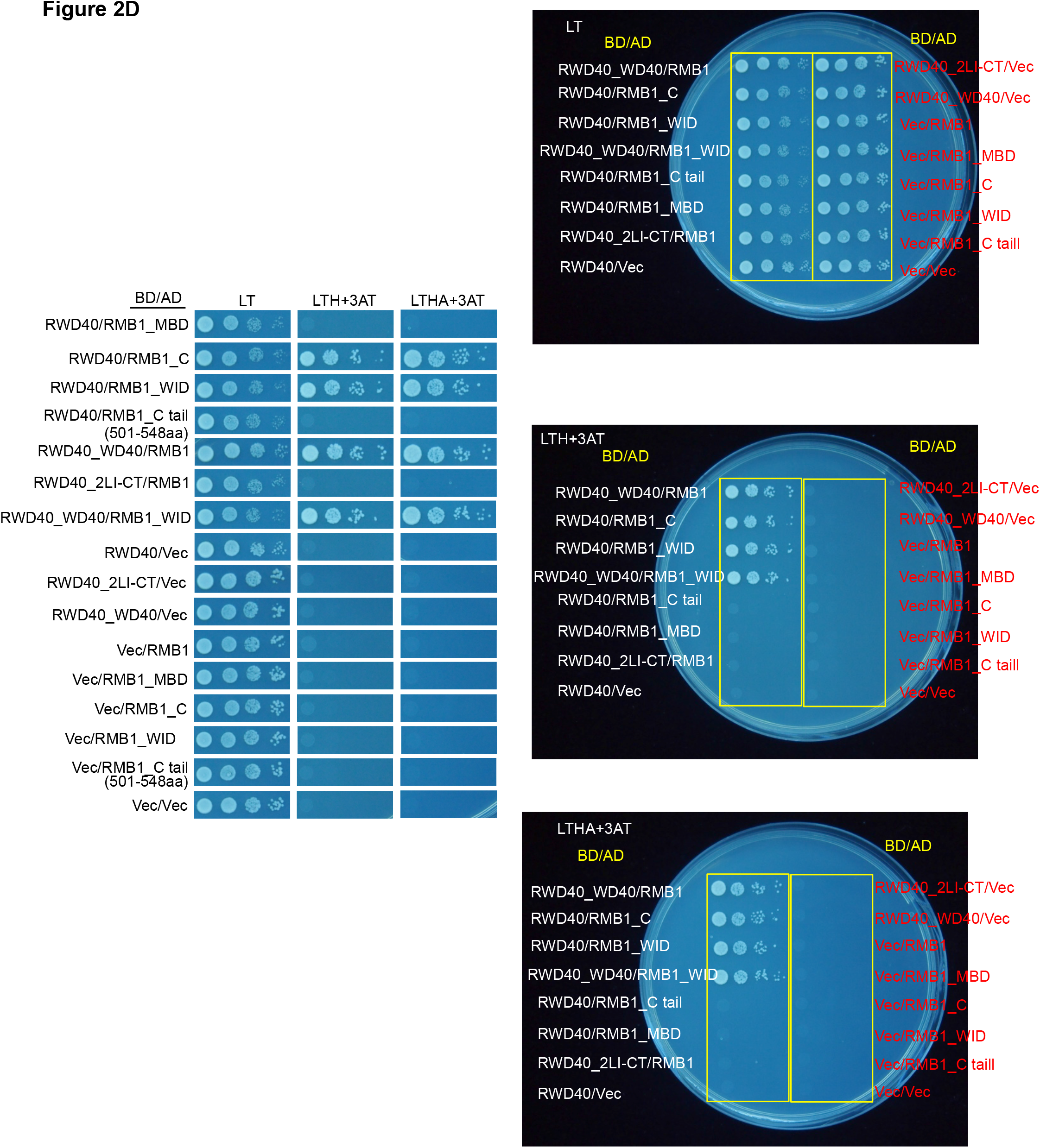
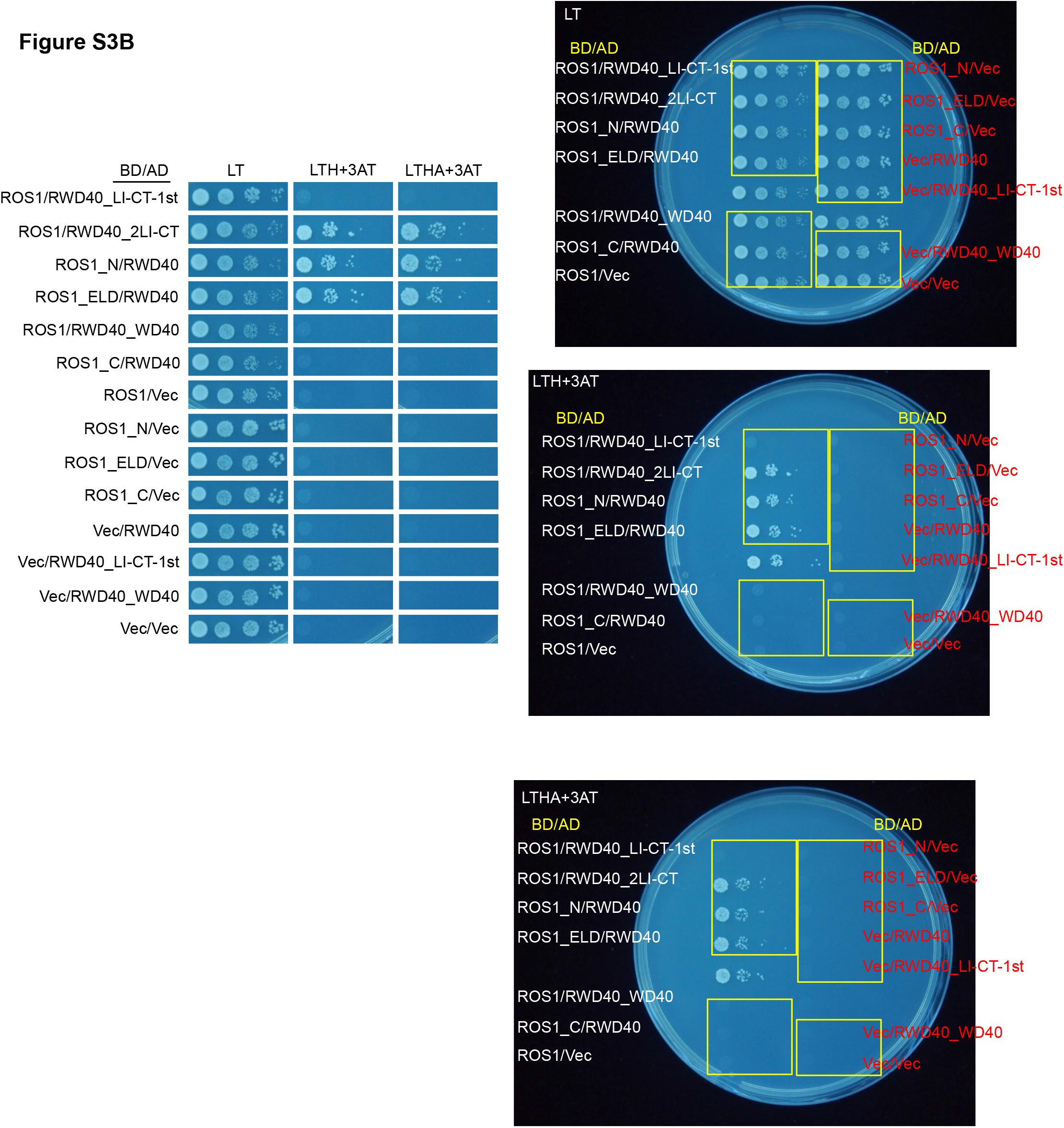
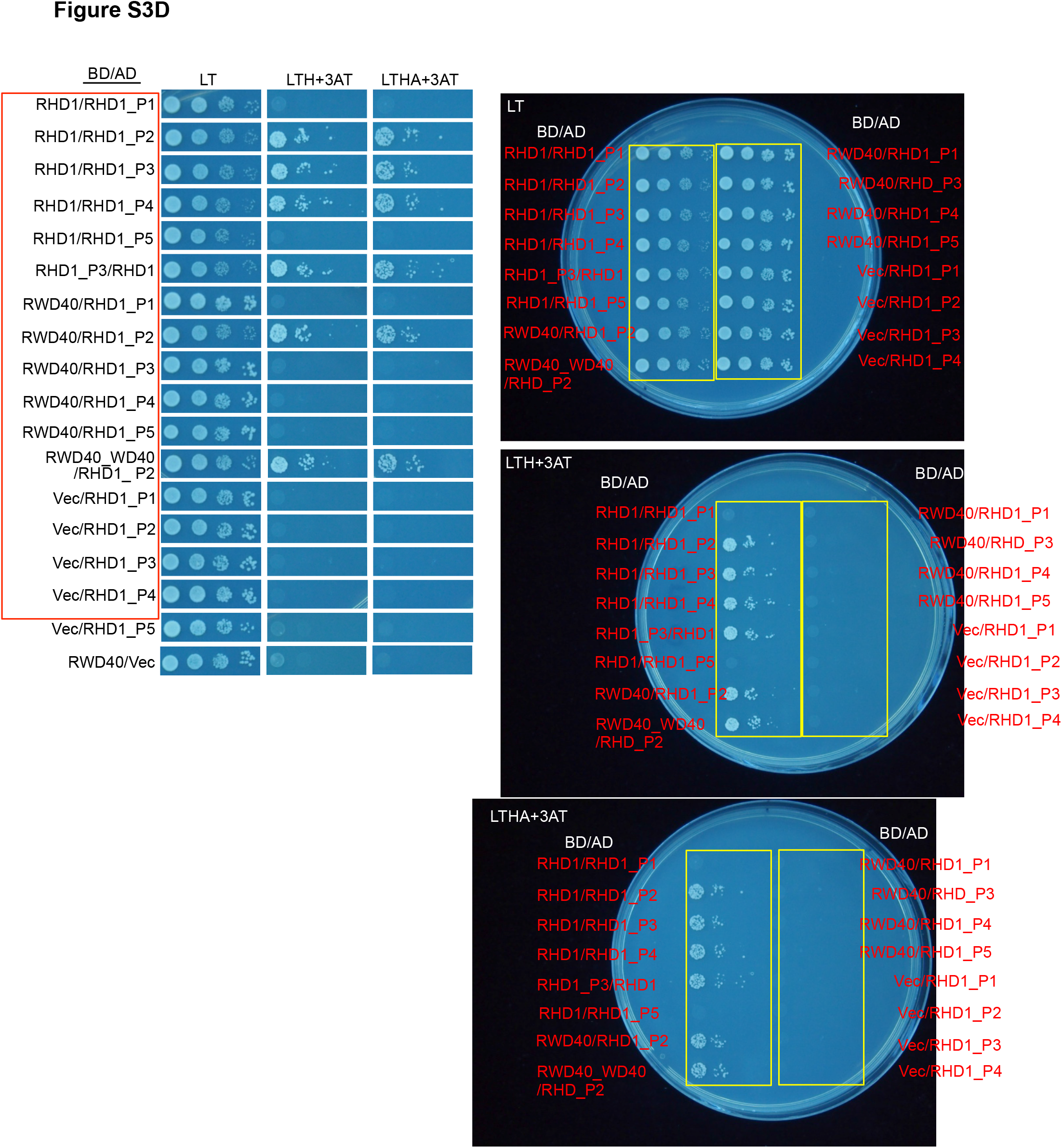
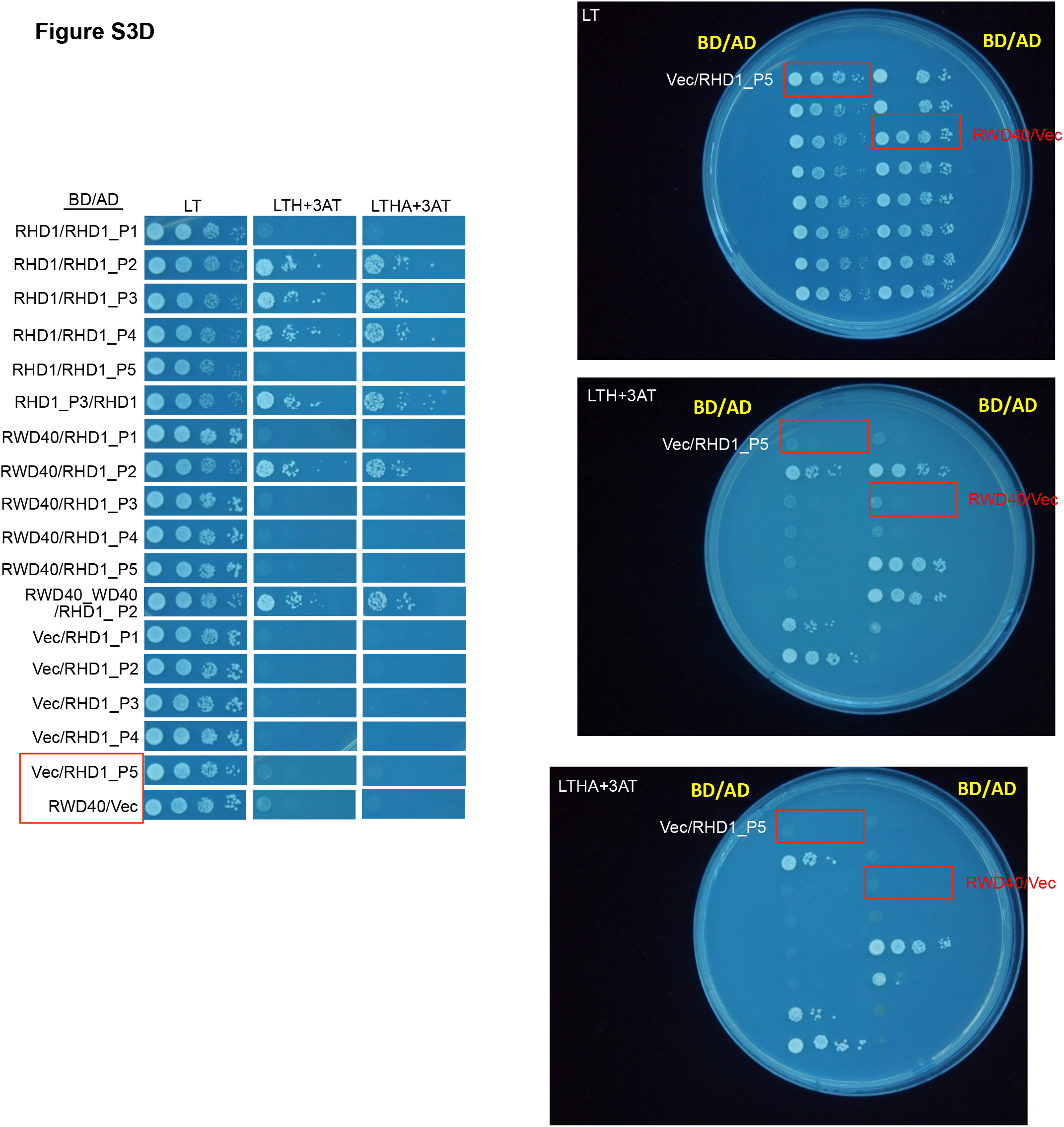
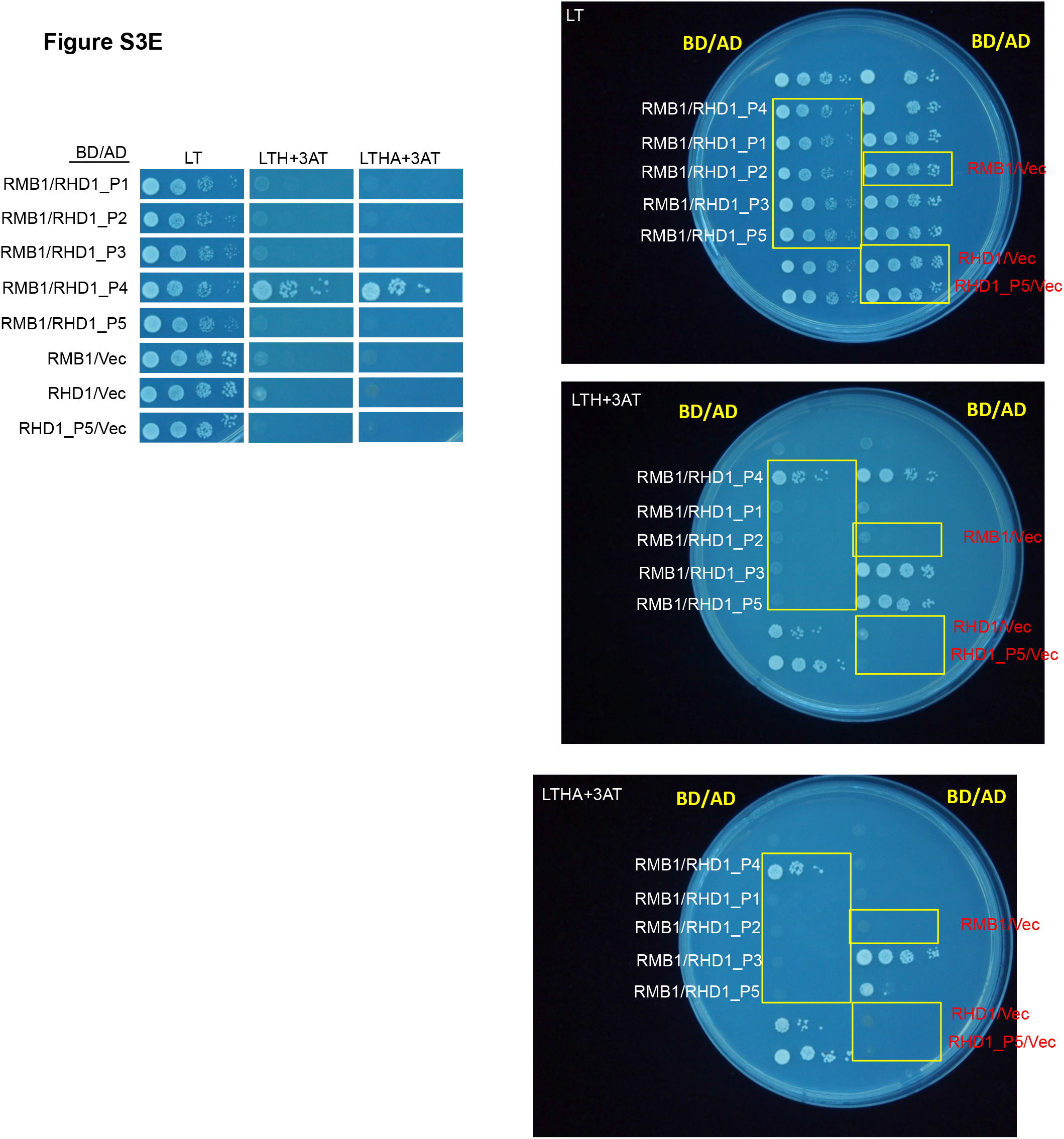
Images of complete Y2H plates (from Figures 1B, 2A, 2D, and S3B-E).

